# Chromosome-scale genome assembly of diploid halophyte *Thinopyrum bessarabicum* excludes J genome from polyploid *Thinopyrum* ancestry

**DOI:** 10.64898/2026.05.20.726494

**Authors:** Nicola Walter, Jack Walker, Cai-yun Yang, Duncan Scholefield, Stephen Ashling, Gemy G. Kaithakottil, David Swarbreck, Aleyda Sierra-Gonzalez, Katie Hawkins, Jonathan A. Atkinson, Darren M. Wells, Malcolm J. Hawkesford, Jianxia Niu, Jesús Quiroz-Chávez, Emile Cavalet-Giora, Simon G. Krattinger, Weilong Guo, Ian P. King, Julie King, Surbhi Grewal

## Abstract

Wild relatives of wheat harbour genetic diversity essential for improving resilience to climate-driven stresses, yet their deployment is hampered by unresolved evolutionary relationships and the absence of reference genomes. Here we present a chromosome-scale reference genome for *Thinopyrum bessarabicum*, a diploid halophyte and high-priority donor for wheat salt tolerance breeding. A key unresolved question is whether the diploid J genome contributed directly to the subgenome composition of extant polyploid *Thinopyrum* species, and which genomic features underpin its exceptional salt tolerance. Using this resource, we show that the diploid J genome of *Th. bessarabicum* is not represented among the subgenomes of polyploid *Thinopyrum* species, resolving a long-standing ambiguity in Triticeae genomics. Gene-level resolution of the reciprocal 4/5 chromosomal translocation across six related Triticeae species identifies conserved breakpoint gene pairs, supporting a single ancestral rearrangement. Genome-wide gene content analysis shows that halophytic capacity is underpinned by quantitative expansion of conserved stress-response gene families. Salt tolerance phenotyping validates chromosome 5J as a tolerance locus in both *Th. bessarabicum* and wheat introgression lines. A physically anchored marker framework and dual-reference skim-sequencing pipeline enable precise megabase-resolution characterisation of *Th. bessarabicum* introgressions in wheat, providing a genomic foundation for deploying J-genome diversity in crop improvement.

## Introduction

Bread wheat (*Triticum aestivum* L.) supplies approximately 20% of global human caloric intake, yet its domestication has left it with a narrow genetic base that constrains its resilience to emerging biotic and abiotic stresses^1, 2^. Wild relatives within the Triticeae tribe represent a vast reservoir of genetic diversity, and their exploitation through introgression has been central to broadening the wheat gene pool^3^. Among these, species of the genus *Thinopyrum* have been extensively used as donors of disease resistance, abiotic stress tolerance, and yield-related traits, particularly in polyploid forms^4–6^.

Despite their widespread use, the genomic composition and evolutionary relationships within *Thinopyrum* remain incompletely resolved. A major source of ambiguity lies in the designation of the “J genome”, which has been applied inconsistently across diploid and polyploid species. The diploid species *Thinopyrum bessarabicum* (Săvul. & Rayss) Á. Löve (2*n* = 2*x* = 14, JJ; E^b^E^b^) is classically defined as carrying a J genome, while the closely related diploid *Thinopyrum elongatum* (Host) D.R. Dewey (2*n* = 2*x* = 14) is assigned an E genome (EE or J^e^J^e^). Although these genomes have often been grouped within a common lineage^7, 8^, other studies have demonstrated that they are genetically distinct and differ in chromosome structure^9–11^, highlighting uncertainty in their evolutionary relationship. *Th. bessarabicum* has nevertheless been widely regarded as a progenitor of several polyploid *Thinopyrum* species under classical cytogenetic models^7, 12–14^.

Polyploid *Thinopyrum* species exhibit complex and often conflicting genomic compositions. Decaploid species such as *Thinopyrum ponticum* (Podp.) Z-W. Liu & R.-C. Wang and *Thinopyrum elongatum* (2 *n* = 10 *x* = 70, EEEEEEE^st^E^st^E^st^E^st^ or JJJJJJJ^st^J^st^J^st^J^st^), carry formulae incorporating multiple J and E-related subgenomes, yet reference genomes for these species remain unavailable due to their size, repetitiveness and structural complexity. Cytogenetic and molecular evidence suggests their genomes are mosaics of subgenomes derived from *Th. bessarabicum* (J), *Th. elongatum* (E)*, Dasypyrum villosum* (L.) P. Candargy (2*n* = 2*x* = 14, VV) and *Pseudoroegneria* lineages (St)^15–18^. The recently published genome of hexaploid *Thinopyrum intermedium* (Host) Barkworth & D. R. Dewey (2*n* = 6*x* = 42), which carries a similarly contested J-containing formula (J^vs^J^vs^J^r^J^r^StSt)^12, 13^, revealed that its three subgenomes derive from *Pseudoroegneria* (St), *Dasypyrum villosum* (L.) P. Candargy (V) and an *Aegilops*-related lineage, not from diploid *Thinopyrum*^19^. However, that analysis included the diploid *Th. elongatum* reference sequence^20^ but not *Th. bessarabicum*, leaving the diploid J genome itself untested against either hexaploid or decaploid *Thinopyrum* lineages. Whether J-designated subgenomes represent diverged derivatives of the diploid J lineage or contributions from a distinct genomic source therefore remained unresolved prior to this study.

*Thinopyrum bessarabicum* is a natural halophyte, capable of surviving at high salinities along the Black Sea coastline^21, 22^, making it one of the most salt-tolerant grasses in the Triticeae and a prioritised donor for wheat salt tolerance breeding^23, 24^. Introgression from *Th. bessarabicum* has contributed genetic variation for a range of agronomically important traits, including disease resistance, abiotic stress tolerance, blue grain and yield-related traits^25–29^. A chromosome-scale reference genome would enable the precise characterisation of germplasm generated through wheat-*Th. bessarabicum* introgression programmes, allowing accurate localisation of introgressed segments and resolution of their gene content, beyond what has been achievable using predominantly cytogenetic approaches^27, 30–33^.

Here, we present the first chromosome-scale, annotated reference genome for *Th. bessarabicum*, generated using PacBio HiFi sequencing and Hi-C scaffolding into seven pseudomolecules. Using this resource, we resolve the evolutionary position of the diploid J genome relative to polyploid *Thinopyrum* species, define the breakpoints of the 4/5 chromosomal translocation at gene-level resolution, and characterise genome-wide patterns of repeat expansion and gene family diversification. We further establish a physically anchored marker framework and a skim-sequencing pipeline enabling high-resolution tracking of *Th. bessarabicum* introgressions in wheat. Together, these results provide a comprehensive genomic foundation for the precise and informed deployment of *Th. bessarabicum* J-genome diversity in wheat improvement programmes.

## Results

### Chromosome-scale assembly and annotation of the *Th. bessarabicum* genome

We generated a chromosome-scale reference genome for *Th. bessarabicum* accession PI531712 using 312 Gb of PacBio HiFi data (∼54× genome coverage) and 662 Gb of Hi-C data (∼112×) (**Supplementary Tables 1-2; Supplementary Fig. 1**). Hifiasm assembly followed by YaHS scaffolding and manual Hi-C curation yielded seven pseudomolecules (Chr1J-Chr7J) anchoring 97.9% of the assembled genome (5.75 Gb; **Fig. 1a**; **Supplementary Table 3**). The final assembly spans 5.87 Gb with a scaffold N50 of 792.8 Mb and a contig N50 of 154.8 Mb (**Table 1**; **Supplementary Table 4**). Chromosome sizes range from 727 Mb (Chr6J) to 946 Mb (Chr2J), with variable contig composition across chromosomes (**Fig. 1b**; **Supplementary Table 3**). Telomeric repeats were identified at five chromosome termini (**Fig. 1a**; black dots) and chloroplast and mitochondrial genomes assembled separately (135,024 bp and 401,710 bp; **Supplementary Table 5**). Genome-wide GC content shows distinct chromosomal patterns, with elevated levels toward pericentromeric regions and lower levels across distal arms (**Fig. 1c**). Assembly accuracy was confirmed by Merqury *k*-mer analysis (QV = 70.85, *k*-mer completeness = 96.77%), Hi-C contact map integrity, BUSCO completeness (98.4% genome mode, 99.3% protein mode against the Poales dataset) and a mean LAI score of 13.86, indicative of reference quality (**Supplementary Fig. 2a-d**).

**Figure 1.**
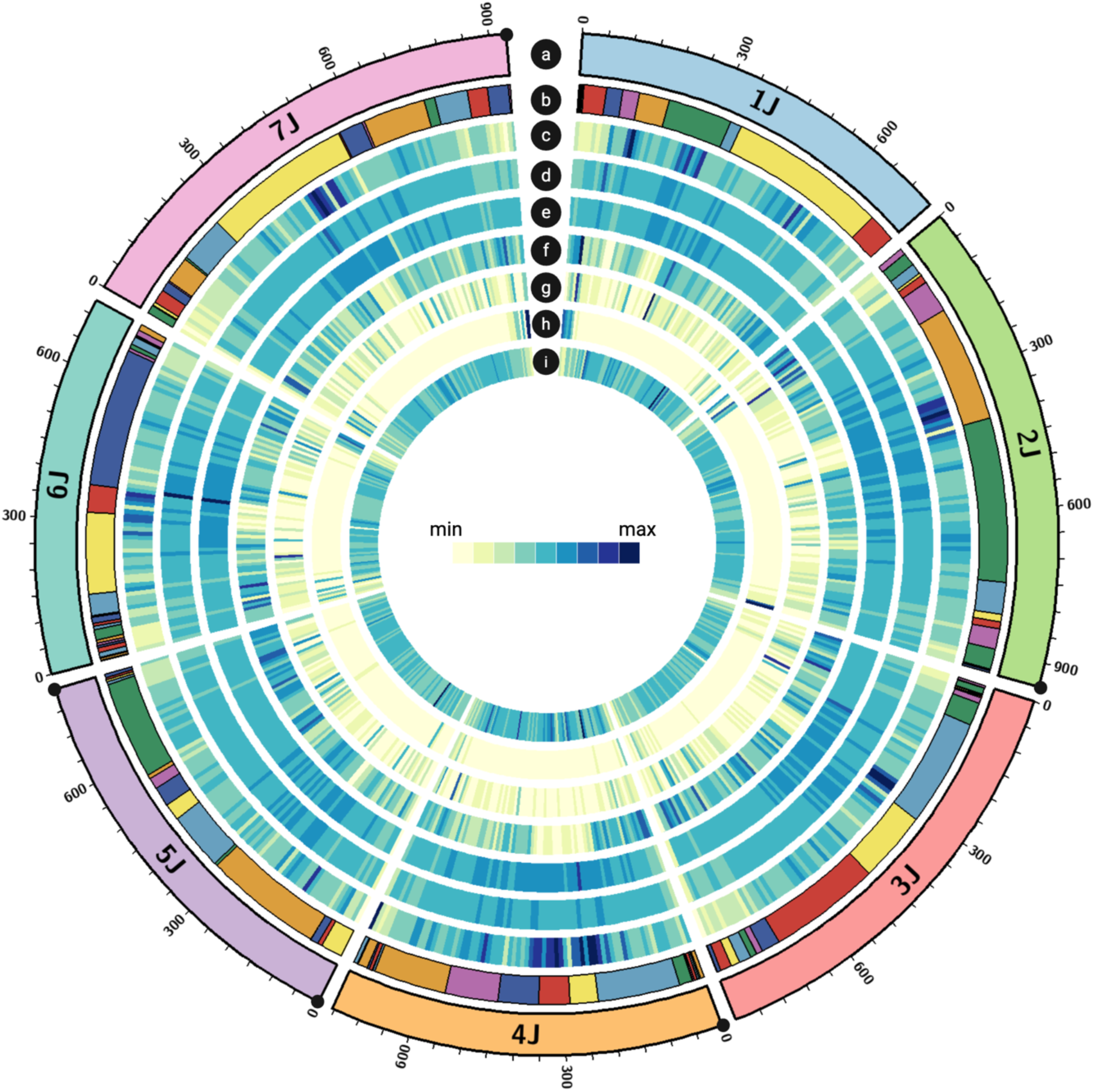
Circos plot showing genomic features across chromosomes 1J–7J. From outer to inner: (**a**) chromosome ideograms with scale (Mb); black dots mark telomeric repeat arrays, (**b**) contig composition of pseudomolecules (minimum resolution 50 Mb), (**c**) GC content, (**d**) total repeat density, **(e)** LTR retrotransposon density, (**f**) high-confidence protein-coding gene density, (**g**) non-coding RNA density, (**h**) nucleotide-binding leucine-rich repeat (NLR) gene density, and (**i)** 5-methylcytosine (5mC) methylation density. All density tracks are calculated in 10 Mb windows. Colours within track B distinguish individual contigs and carry no additional biological meaning.

**Table 1.** Assembly and annotation statistics for the *Th. bessarabicum* nuclear genome.

| **Metric** | **Value** |
| --- | --- |
| Assembly size (bp) | 5,873,390,317 |
| Number of scaffolds | 502 |
| Number of contigs | 646 |
| Scaffold N50 (bp) | 792,792,867 |
| Scaffold L50 | 4 |
| Contig N50 (bp) | 154,753,986 |
| Contig L50 | 12 |
| Largest scaffold (bp) | 946,002,307 |
| Anchored size (bp) | 5,750,579,165 |
| GC content (%) | 45.78 |
| Number of gaps | 144 |
| **Feature (high-confidence protein-coding)** | **Number or length** |
| Number of genes | 36,453 |
| Number of transcripts | 66,267 |
| Transcripts per gene | 1.82 |
| Maximum gene length (bp) | 74,474 |
| Minimum gene length (bp) | 96 |
| Average gene length (bp) | 3,506 |
| Average protein length (aa) | 403 |
| Average 5' UTR length (bp) | 162 |
| Average 3' UTR length (bp) | 298 |
| Exons per gene | 4.54 |
| Average exon length (bp) | 367 |
| Average intron length (bp) | 518 |

Repeat annotation identified 80.6% of the genome as repetitive DNA (**Fig. 1d-e**; **Supplementary Table 6**). Evidence-based annotation integrating RNA-Seq and Iso-Seq transcriptomic data from six tissue types (**Supplementary Tables 7-8**), cross-species protein alignments from ten grass proteomes (**Supplementary Table 9**), and *ab initio* prediction yielded 114,793 gene models across all biotype categories, corresponding to 150,167 transcript isoforms (**Fig. 1f-g; Supplementary Tables 10 -11**). Of 89,641 protein-coding gene models, 36,453 were designated high-confidence (HC), with 98.7% functionally annotated (**Table 1**; **Supplementary Table 11**). An additional 16,367 transposable element-related gene models were identified, of which 2,274 were classified as high-confidence (**Supplementary Table 11**). NLR resistance genes showed distal chromosomal enrichment, particularly on chromosomes 2J, 3J and 7J, mirroring patterns seen in other Triticeae assemblies (**Fig. 1h**; **Supplementary Table 12**). Other defence-associated gene families, including receptor-like kinases, ABC transporters, cytochrome P450s and WRKY transcription factors, show similar clustered distributions, particularly on chromosomes 2J, 3J and 7J (**Supplementary Fig. 3a-b; Supplementary Table 12**). Genome-wide 5mC methylation density showed pronounced enrichment across repeat-rich pericentromeric regions, contrasting with the distal enrichment of protein-coding genes and NLR loci (**Fig. 1i**).

### Evolutionary divergence of the diploid J genome from polyploid Thinopyrum subgenomes

OrthoFinder assigned 97.2% of genes to 64,307 orthogroups across 29 species. Of 1,539 orthogroups shared by all species, 210 single-copy orthologues were retained for phylogenomic inference. Maximum-likelihood and multispecies coalescent analyses produced congruent, strongly supported topologies (**Fig. 2a**). *Th. bessarabicum* clustered robustly with diploid *Th. elongatum* as a distinct J/E lineage, clearly separated from all *Th. intermedium* subgenomes: the *Th. intermedium* “J” subgenome clustered with D-genome species, and the remaining subgenomes showed affinities to *Pseudoroegneria* (St) and *Dasypyrum* (V) lineages. Divergence time estimation placed the split between the diploid J/E lineage and the broader Triticeae clade at ∼6.7 Mya (95% HPD: 5.6-8.0 Mya; **Fig. 2a**), substantially predating the origin of polyploid *Th. intermedium*.

**Figure 2.**
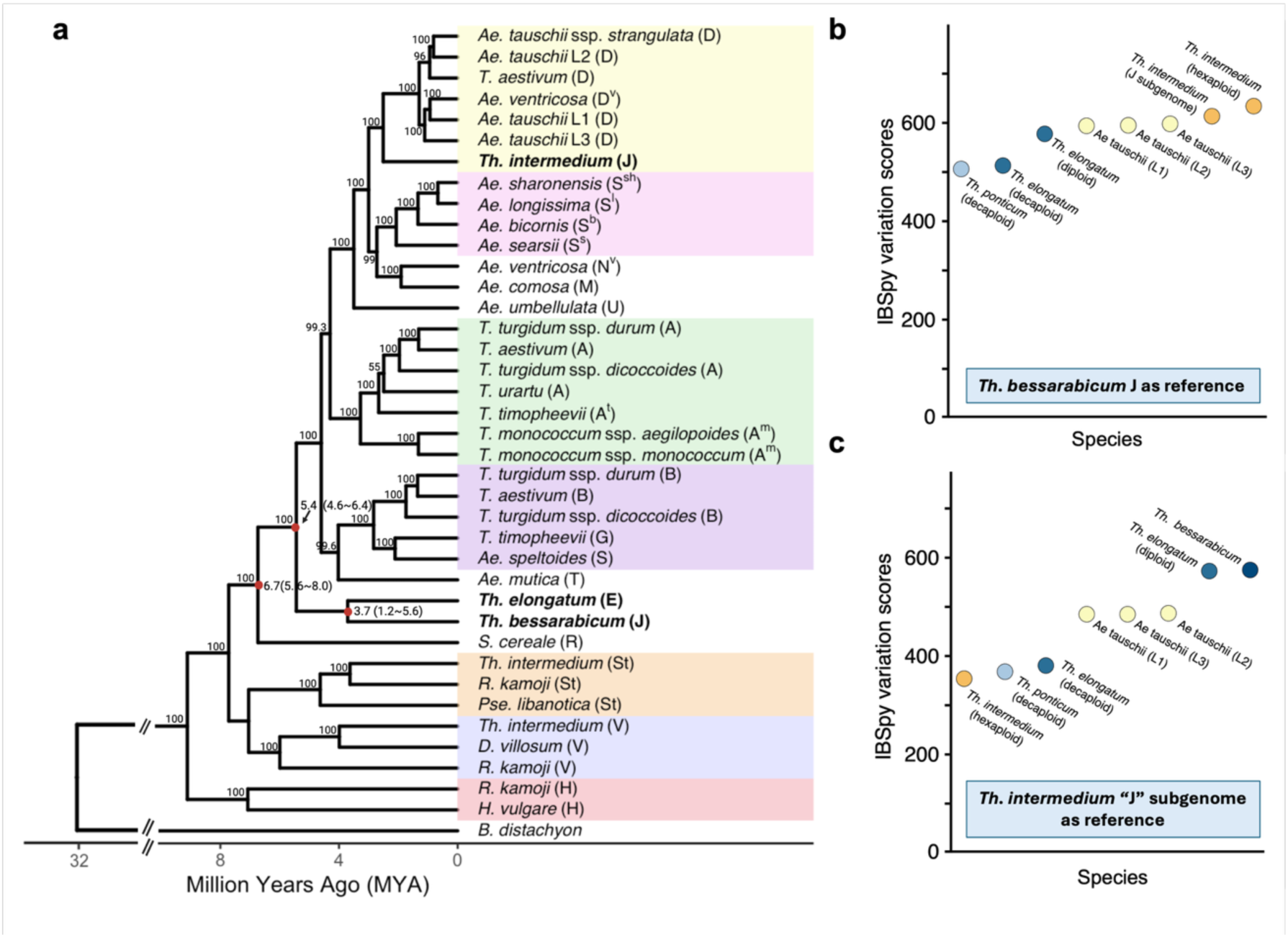
Phylogenomic and *k*-mer–based evidence for divergence of the diploid J genome from polyploid *Thinopyrum* subgenomes. (a) Time-calibrated maximum-likelihood phylogeny of representative Triticeae species inferred from 210 single-copy orthologous genes. Numbers at nodes indicate maximum-likelihood bootstrap support values based on 1,000 replicates. Red circles mark selected evolutionary nodes for which divergence times and 95% highest posterior density (HPD) intervals are shown. Shaded boxes denote major genomic lineages (A, B, D, St, V, S and H). (b) Comparison of genome similarity using IBSpy variation scores calculated from *k*-mer sets aligned to the diploid *Th. bessarabicum* J genome reference. *k*-mer sets were derived from the whole-genome sequence of hexaploid *Th. intermedium* and its assembled “J” subgenome independently, as well as from decaploid *Th. elongatum*, decaploid *Th. ponticum*, diploid *Th. elongatum*, and three *Ae. tauschii* lineages (L1-3). All polyploid *Thinopyrum* accessions, including both the whole-genome and J-subgenome *k*-mer sets of *Th. intermedium*, display variation scores exceeding 500, indicating different-species divergence from the diploid J genome under established thresholds. (**c**) IBSpy variation scores calculated using the *Th. intermedium* “J” subgenome as the reference. Polyploid *Thinopyrum* species exhibit intermediate variation scores (∼300–400), consistent with divergence within a shared genomic lineage.

Assembly-free *k*-mer analysis using IBSpy corroborated this result. When the *Th. bessarabicum* J genome was used as reference, all polyploid accessions tested, decaploid *Th. elongatum* and *Th. ponticum* and hexaploid *Th. intermedium*, displayed IBSpy variation scores exceeding 500 across all seven chromosomes (**Fig. 2b**; **Supplementary Fig. 4**). Based on identity thresholds defined for *Ae. tauschii* IBSpy varation scores >500 indicate different species^34, 35^. When the *Th. intermedium* "J" subgenome was used as reference, polyploid *Thinopyrum* accessions showed lower variation scores (∼300-400; **Fig. 2c**) indicative of within-species diversification. Notably, *Ae. tauschii* accessions representing the D lineage showed higher variation scores than polyploid *Thinopyrum* relative to the *Th. intermedium* "J" reference yet remained below the scores observed for the diploid J/E lineage. This ordering is consistent with the phylogenomic placement of the *Th. intermedium* "J" subgenome within the D-genome clade (**Fig. 2a**) and confirms that the intermediate scores of polyploid *Thinopyrum* reflect shared ancestry with the *Th. intermedium* "J" lineage rather than with the diploid J genome of *Th. bessarabicum*. Together, these analyses demonstrate that the diploid J genome of *Th. bessarabicum* is not represented among the subgenomes of extant polyploid *Thinopyrum* species.

### Chromosome structural rearrangements and comparative synteny

Pairwise gene-based synteny shows that the J genome carries a reciprocal translocation between homoeologous chromosomes 4 and 5, absent in *Th. elongatum* (E), *Th. intermedium* (“J” subgenome) and *Aegilops tauschii* but shared with *D. villosum* (V), *Triticum urartu* (A) and bread wheat (**Supplementary Fig. 5**). *Th. bessarabicum* and *Th. elongatum* showed extensive collinearity across all seven chromosome pairs, with the sole exception of the distal arms of chromosomes 4J and 5J (**Supplementary Fig. 5a**): the distal region of Chr4J (663 to ∼752 Mb; 2,369 genes) is collinear with Chr5E, while the distal region of Chr5J (712 to 792 Mb; 2,081 genes) is collinear with Chr4E, confirming the reciprocal nature of the rearrangement.

To place the rearrangement in its broader evolutionary context, a six-species comparative karyotype was constructed for homoeologous groups 4 and 5, using *Th. elongatum* as the ancestral unrearranged reference (**Fig. 3a**). *Th. bessarabicum*, *D. villosum* and *T. urartu* all display identical translocated ribbon patterns, with the proximal boundaries of both translocated segments conserved across all three species, while *Th. intermedium* ("J" subgenome), *Ae. tauschii* and *Th. elongatum* retain the ancestral arrangement. *D. villosum* additionally carries a 4/7 translocation absent from both the J and E genomes (**Fig. 3a**; **Supplementary Fig. 5d**), confirming this rearrangement as V-lineage specific and post-dating the shared 4/5 event.

**Figure 3.**
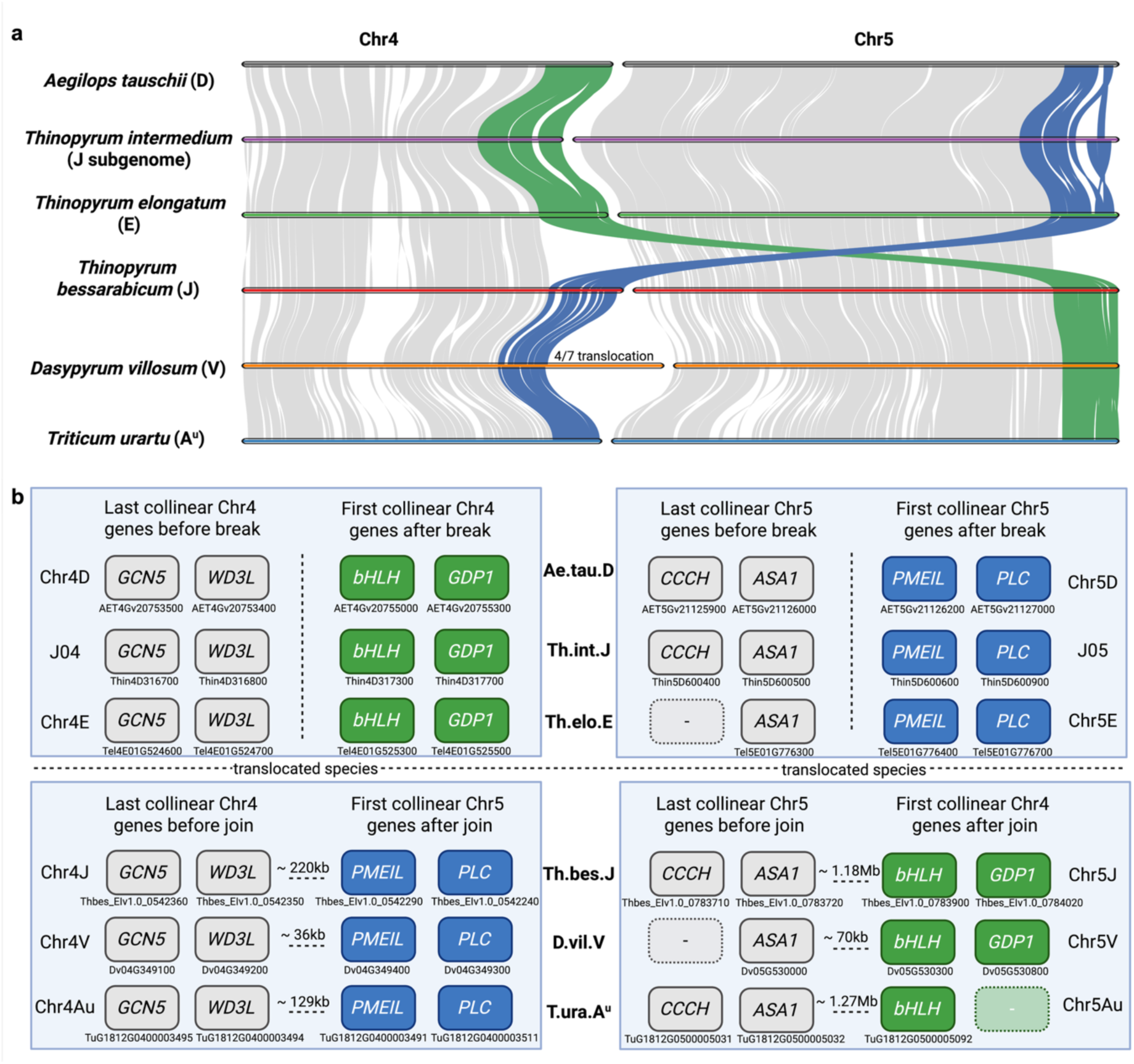
Comparative synteny and microsynteny of the 4/5 chromosomal translocation across six Triticeae species. **(a)** Gene-based karyotype ribbons for homoeologous chromosomes 4 and 5 across *Aegilops tauschii* (D), *Thinopyrum intermedium* “J” subgenome, *Th. elongatum* (E), *Th. bessarabicum* (J), *Dasypyrum villosum* (V) and *Triticum urartu* (A^u^). Grey ribbons indicate collinear syntenic blocks. Green ribbons trace the segment ancestrally derived from the distal Chr4 long arm; blue ribbons trace the reciprocal Chr5 long arm segment. In species retaining the ancestral arrangement (*Ae. tauschii*, *Th. intermedium*, *Th. elongatum*), coloured ribbons connect homoeologous regions on the same chromosome. In species carrying the translocation (*Th. bessarabicum*, *D. villosum*, *T. urartu*), ribbons cross between chromosomes, demonstrating the reciprocal nature of the rearrangement. The additional 4/7 translocation unique to *D. villosum* is annotated. **(b)** Microsynteny at the two chromosomal junction points defining the 4/5 translocation. Left panels show the Chr4 junction; right panels show the Chr5 junction. Upper rows show ancestral species (*Ae. tauschii*, *Th. intermedium* “J” subgenome, *Th. elongatum*); lower rows show translocated species (*Th. bessarabicum*, *D. villosum*, *T. urartu*). Grey boxes: collinear flanking genes. Coloured boxes indicate genes at the breakpoint: green marks Chr4-origin genes, blue marks Chr5-origin genes. Dashed boxes indicate positions where no ortholog was detected. Intergenic distances between the last pre-join and first post-join gene pairs are shown in kilobases or megabases in species with the 4/5 translocation. Gene abbreviations: *GCN5*, GCN5-related N-acetyltransferase 8; *WD3L*, WD repeat-containing protein 3; *bHLH*, basic helix-loop-helix domain-containing protein; *GDP1*, G-patch domain-containing protein 1; *CCCH*, zinc finger CCCH domain-containing protein; *ASA1*, anthranilate synthase alpha subunit 1; *PMEIL*, pectinesterase inhibitor domain-containing protein; *PLC*, phospholipase C. Gene IDs are shown below each box. Full UniProt annotations were confirmed by BLASTp against the UniProt/Swiss-Prot database.

Microsynteny analysis resolved the two translocation breakpoints at gene-pair resolution across all six species (**Fig. 3b**; **Supplementary Table 13**): on Chr4J, the junction occurs between *WD3L* (Chr4J: 662,851,139) and *PMEIL1* (Chr4J: 663,071,327); on Chr5J, between *ASA1* (Chr5J: 711,009,294) and *bHLH* (Chr5J: 712,197,509). Flanking synteny markers including GCN5, GDP1, CCCH-type zinc finger, and PLC genes on either side of the breakpoint confirm collinearity with the homoeologous chromosomes in the non-translocated species (**Fig. 3b**). The orthologous breakpoint gene pairs are identical to those reported in *D. villosum*^36^ and conserved across all tested genomes, pointing to a single ancestral rearrangement predating the divergence of the J, V and A genome lineages. Translocated segments in the J genome are 3.3-fold larger (171.8 Mb vs 52.6 Mb in V) and contain 3.3-fold more genes (4,450 vs 1,356) than their counterparts in *D. villosum* (**Supplementary Table 13**).

### Transposable element dynamics and genome expansion

J genome chromosomes are uniformly larger than those of *Th. elongatum* (E), *Th. intermedium* (“J” subgenome), *D. villosum* (V) and *T. urartu* (A^u^) across all homoeologous groups, with the greatest size difference relative to *Th. elongatum* (mean 1.27-fold; 5.87 Gb vs 4.54 Gb; **Fig. 4a**). To investigate the molecular basis of this size difference, we characterised the transposable element landscape using structural and temporal analyses. Identification of 92,588 intact LTR retrotransposons in the J genome, collectively spanning 904.6 Mb (15.4% of genome; **Supplementary Table 14**), revealed that 89.6% inserted within the last 2 Mya (mean insertion age 1.07 Mya), indicating a pronounced recent transposition burst. In contrast, only 11.2% of intact E genome LTR elements show recent insertion **(Supplementary Table 14)**, representing an 8.0-fold difference in recent transposition activity between the two genomes. Gypsy and Copia elements account for 46.0% and 32.6% of intact J genome LTRs respectively, both showing strong recent activity (87.1% and 95.4% of insertions ≤2 Mya). Subfamily-level classification assigned 47,313 elements (51.1%) to known LTR subfamilies; 45,275 elements (48.9%) remained unclassified despite comprehensive database searches, suggesting the presence of J-genome-specific TE lineages. Among classified subfamilies, ANGELA (Copia; 11,662 copies, 12.6%), SABRINA (Gypsy; 6,061 copies, 6.55%), DERAMI, FATIMA, WHAM and WILMA were the six most abundant, all showing pronounced recent proliferation, with 94.2% of ANGELA and 89.2% of SABRINA insertions within the last 2 Mya (mean insertion ages 1.08 and 1.49 Mya respectively; **Fig. 4b**; **Supplementary Table 15**).

**Figure 4.**
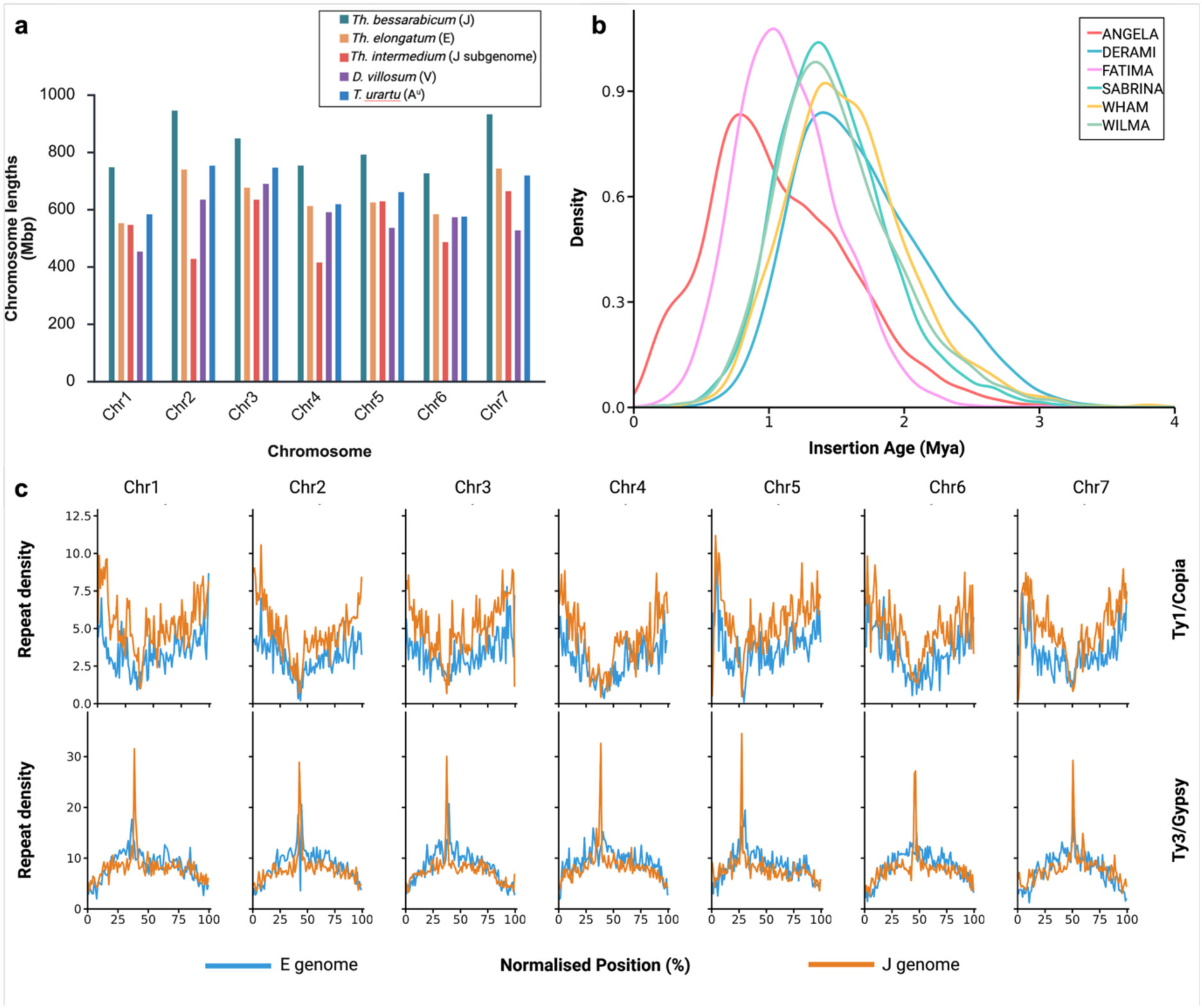
Transposable element dynamics and genome expansion in *Thinopyrum bessarabicum*. **(a)** Chromosome length comparison across homoeologous groups 1-7 for *Th. bessarabicum* (J), *Th. elongatum* (E), *Th. intermedium* (“J” subgenome), *Dasypyrum villosum* (V), and *Triticum urartu* (A^u^). **(b)** Kernel density estimates of insertion age distributions for the six most abundant intact LTR-retrotransposon subfamilies in the *Th. bessarabicum* J genome. **(c)** Density of intact Ty1/Copia and Ty3/Gypsy LTR-retrotransposons along the J and E genome chromosomes. Each chromosome assembly was divided into 100 equal-proportion positional bins and repeat density per bin is expressed as a percentage of bin length, normalising for differences in chromosome length between the J and E genomes.

Comparison with the E genome revealed distinct superfamily dynamics underlying J genome expansion (**Fig. 4c**; **Supplementary Table 14**). Copia and Gypsy elements showed fundamentally different distributional patterns when repeat density was normalised by bin length to account for differences in chromosome length between the J and E genomes. Copia content was consistently and substantially elevated across all seven J genome chromosomes (total: 287.3 Mb vs 151.6 Mb, 1.89-fold genome-wide), with per-chromosome ratios ranging from 1.75 to 2.12-fold and mean Copia density 1.49-fold higher in J than E across all chromosomes. This excess was uniformly distributed along chromosome arms rather than concentrated at any particular chromosomal region, indicating broad genome-wide proliferation of Copia elements. Gypsy showed a contrasting pattern: despite modestly greater total content in J (459.1 Mb vs 389.3 Mb, 1.17-fold), genome-wide mean Gypsy density was comparable between the two genomes (J: 7.95%, E: 8.60% of bin length), with per-chromosome J/E density ratios close to 1.0 across all seven chromosomes (range 0.88-1.01). Instead, Gypsy accumulation in the J genome was strongly centromere-concentrated on all seven chromosomes, with peak centromeric density reaching 34.5% of local sequence in J compared with a genome-wide maximum of 20.7% in the E genome. While a centromeric Gypsy peak was detectable on most E genome chromosomes, its amplitude was variable and markedly lower than in J, with arm-level densities broadly equivalent between the two genomes. Centromeric domains were delimited using smoothed CEREBA-lineage LTR density, defining compact domains on all seven chromosomes ranging from 8.2 Mb (Chr1J) to 11.5 Mb (Chr2J), mean 9.8 Mb, cross-validated against gene-poor regions (**Supplementary Fig. 6; Supplementary Table 16**).

### Comparative genomics of salt-tolerance gene content

Salt-tolerance gene content was analysed across three orthogroup sets: 58 *Th. bessarabicum*-specific orthogroups (83 HC genes), 201 orthogroups shared across *Thinopyrum* species but absent from bread wheat, and 20,078 orthogroups shared with bread wheat (35,170 HC genes; **Supplementary Fig. 7; Supplementary Tables 17 -19**). Among J-genome-specific genes, 18 genes from 5 orthogroups carry salt-relevant annotation including salt-induced protein 3 and dehydrins, with no enrichment above genome-wide background (**Supplementary Table 17**). Among *Thinopyrum*-shared, wheat-absent orthogroups, 108 salt-relevant genes from 26 orthogroups were identified (41 HC; **Supplementary Table 18**), including LEA-2 proteins, AP2/ERF transcription factors, germin-like proteins, and the most prominent family OG0000623. This NAC domain transcription factor orthogroup is absent from all three bread wheat subgenomes yet carries 57 copies in *Th. bessarabicum* (10 HC, 47 LC), concentrated in a 6.34 Mb tandem array at 706-712 Mb on Chr4J, compared to 28 copies across *Th. intermedium* subgenomes, 15 in *D. villosum*, 7 in *H. vulgare*, and 1 in *Th. elongatum* (**Supplementary Fig. 8**).

Across the 20,078 shared orthogroups, 14 of 15 salt-relevant gene family categories showed significantly higher mean copy numbers in *Th. bessarabicum* than in bread wheat (**Supplementary Fig. 9a; Supplementary Table 19**), with NAC and DREB/AP2/ERF families showing the greatest enrichment (8.06-fold and 5.05-fold respectively; both p < 0.001). Chromosomal mapping of HC salt-gene density and copy number enrichment revealed broad genome-wide enrichment, with a pronounced cluster of both high gene density and elevated copy number ratios on the distal Chr5J long arm at 758-780 Mb (**Supplementary Fig. 9b**). The genome-wide distribution of salt-relevant HC genes along Chr5J (**Fig. 5a**) shows that this hotspot sits within a broader region of elevated AP2/ERF and NAC gene density on the long arm, with an AP2/ERF cluster at approximately 496 Mb comprising 40 HC genes from 14 evolutionarily distinct orthogroups, each independently expanded from a different ancestral subfamily, and two NAC domain transcription factor arrays at approximately 626 Mb and 725 Mb. Detailed examination of the 758-780 Mb hotspot (**Fig. 5b-c**) revealed orthogroup OG0000191 forming a 52-gene HC germin-like protein (GLP) tandem array spanning 0.69 Mb at approximately 772-773 Mb, carrying 8.84-fold more gene copies genome-wide than bread wheat. Co-localised at this hotspot are representatives of two LEA-2 orthogroups, a bZIP transcription factor orthogroup and a sodium transporter orthogroup, each showing elevated genome-wide copy numbers relative to bread wheat (24-fold, 8-fold, 4-fold and 2-fold respectively; **Fig. 5b**). Across the full GLP complement, *Th. bessarabicum* carries 156 HC GLP copies distributed genome-wide, with the highest OG0000191 copy number of any of the 38 Triticeae species in the panel, while *Th. intermedium* carries only 3 copies of this orthogroup, indicating J-genome-specific amplification (**Supplementary Fig. 10**).

**Figure 5.**
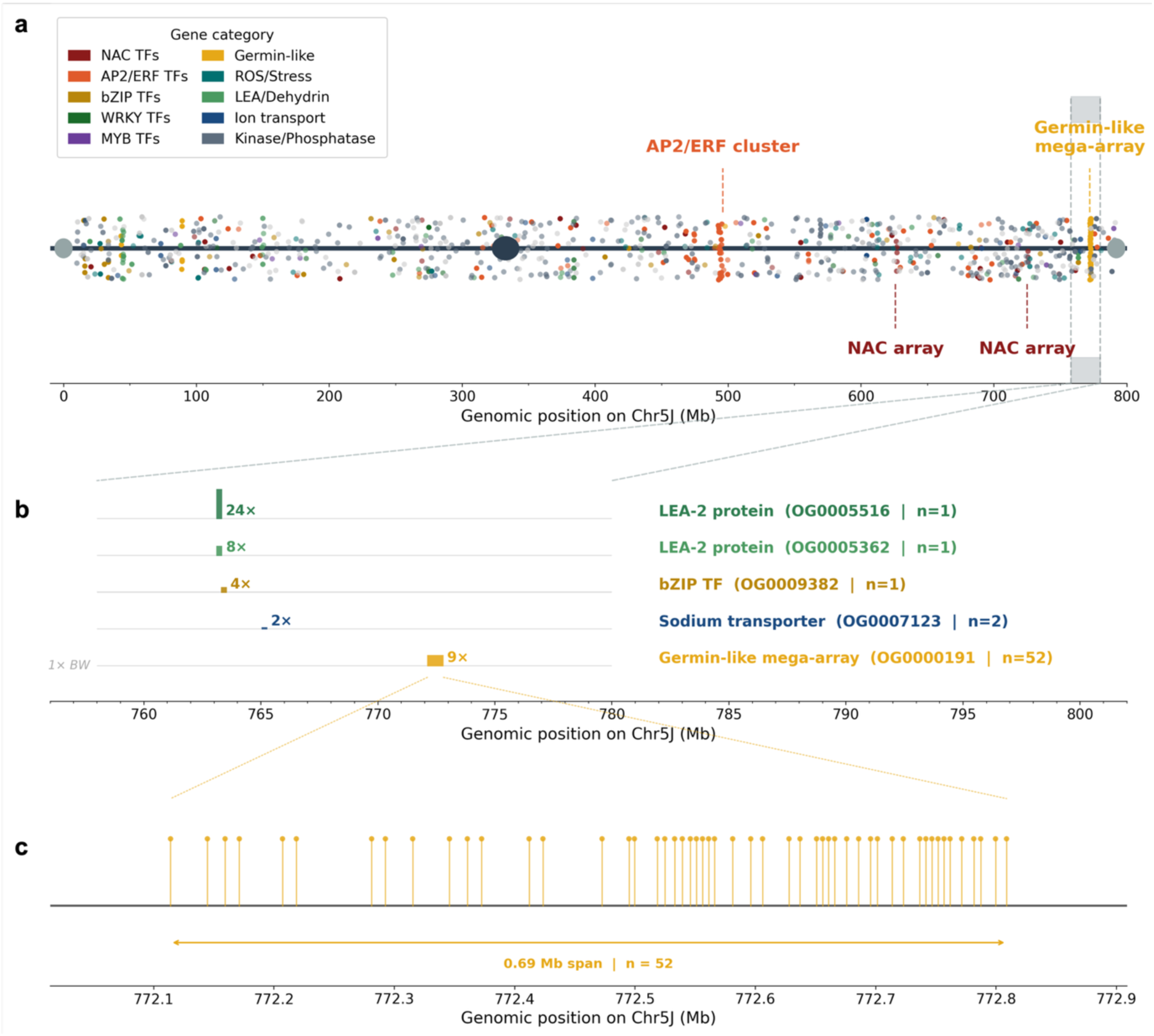
Salt-relevant candidate gene landscape of Chr5J in *Th. bessarabicum*. **(a)** Distribution of high-confidence (HC) salt-relevant candidate genes across Chr5J. Each dot represents one HC gene coloured by functional category; dot opacity is scaled to the *Th. bessarabicum*/bread wheat copy number ratio (brighter = higher ratio). The centromere is indicated by the filled ellipse. Vertical dashed lines and grey shading mark the 758 to 780 Mb region expanded in panel B. Notable clusters are annotated: an AP2/ERF transcription factor cluster at approximately 496 Mb, two NAC domain transcription factor arrays at approximately 626 Mb and 725 Mb, and the germin-like protein mega-array at approximately 772 Mb. **(b)** Zoom of the 758 to 780 Mb stress-gene hotspot. Each horizontal track shows the genomic span and relative copy number of one orthogroup. Bar height is proportional to the *Th. bessarabicum*/bread wheat copy number ratio; the grey baseline represents the bread wheat reference level (1x). Orthogroup identifier and the number of *Th. bessarabicum* HC gene copies (n) are shown to the right of each track. **(c)** Genomic organisation of the OG0000191 tandem array on Chr5J. Each vertical line and filled circle represents one HC gene model. The array comprises 52 HC genes spanning 0.69 Mb at approximately 772 Mb on Chr5J.

### Genomic characterisation and salt tolerance phenotyping of wheat-*Th. bessarabicum* **introgression lines**

Physical mapping of 277 KASP markers to the J genome assembly resolved their distribution across all seven J chromosomes at a median inter-marker spacing of 11.6 Mb (**Supplementary Table 20; Supplementary Fig. 11**), anchoring the complete panel to J-genome chromosomal coordinates for the first time. Of these, 170 markers were originally validated against *Th. bessarabicum*^37^; the remaining 107, developed for other wild relative species^38, 39^, are reported here as also polymorphic between wheat and *Th. bessarabicum*. Using this panel for marker-assisted selection during backcrossing, five homozygous wheat-*Th. bessarabicum* introgression lines (WRC22-Bess1-5) were generated and are reported here for the first time. Dual-reference skim-sequencing against the bread wheat RefSeq v2.1^40^ and the *Th. bessarabicum* assembly confirmed J chromosome of origin and introgression boundaries at 1 Mb resolution for all five lines (**Supplementary Table 21**), including one unusual disomic addition chromosome comprising joined segments of Chr1J and Chr6J with an additional A-genome fragment confirmed by GISH (**Supplementary Figure S12**).

The same dual-reference pipeline was applied to 20 genebank wheat-*Th. bessarabicum* lines including disomic additions, substitution lines, centric fusions and translocations. Three lines previously designated as simple alien additions were confirmed as centric fusions between two J chromosomes^30^, with simultaneous gain on both contributing J chromosomes (**Supplementary Fig. 13a-c**), and one line (T5AS.5JL) was confirmed as a Robertsonian translocation between Chr5J and wheat chromosome 5A (**Supplementary Fig. 13d**). Translocation line T4BS.4BL-4JL, characterised by Patokar et al. (2016)^31^ as carrying a Chr4JL segment, instead shows Chr5J gain, demonstrating that the introgressed segment originates from Chr5J (**Supplementary Table 21**).

To validate the functional relevance of the Chr5J salt tolerance gene content identified above, and to characterise the *Th. bessarabicum* accessions from which the introgression lines were derived, salt tolerance was evaluated phenotypically in two independent experiments. The genome assembly reported here was generated from *Th. bessarabicum* accession PI 531712. This accession failed to establish with sufficient replication under controlled conditions due to poor seed germination, precluding its direct inclusion in phenotypic analyses. Germination screening was therefore conducted on accessions PI 531711 and PI 8388607, two further *Th. bessarabicum* accessions available within the germplasm collection. Paragon and Shiraz were included as bread wheat controls to provide a representative baseline of germination-stage performance in *T. aestivum*. Chinese Spring (CS) and Wembley were used as controls in the hydroponic experiment, where CS is the genetic background of the introgression lines and Wembley is an established salt-sensitive reference.

At germination stage, significant genotypic variation in root length and shoot length salt tolerance index (RLSTI and SLSTI respectively) was detected across NaCl concentrations (Kruskal-Wallis, p < 0.05 at all stress concentrations). Representative seedlings illustrating the progressive reduction in root and shoot growth under increasing NaCl concentrations are shown for wheat cultivars and *Th. bessarabicum* accessions (**Fig. 6a**). Both *Th. bessarabicum* accessions maintained greater RLSTI than wheat cultivars under progressive salt stress (**Fig. 6b**). PI 531711 was significantly more tolerant than Shiraz at 100 mM NaCl (p < 0.05) and significantly more tolerant than both wheat cultivars at 150 and 250 mM (p < 0.05), retaining 107.9% and 67.4% of its unstressed root length respectively at these concentrations. At 150 mM NaCl, PI 531711 RLSTI exceeded 100%, indicating that root growth was maintained above unstressed control levels at this concentration. PI 8388607 showed significant tolerance vs Shiraz at 100 mM and vs both cultivars at 150 mM (p < 0.05), though at 250 mM only two replicate seedlings established, precluding statistical comparison. For SLSTI (**Fig. 6c**), PI 531711 significantly outperformed both wheat cultivars at 150 mM (p < 0.05) and Shiraz at 250 mM (p < 0.05), with shoot growth maintained at 125% of the unstressed control at 150 mM. PI 8388607 showed a trend toward improved SLSTI over wheat at most concentrations, though differences did not reach significance after correction.

**Figure 6.**
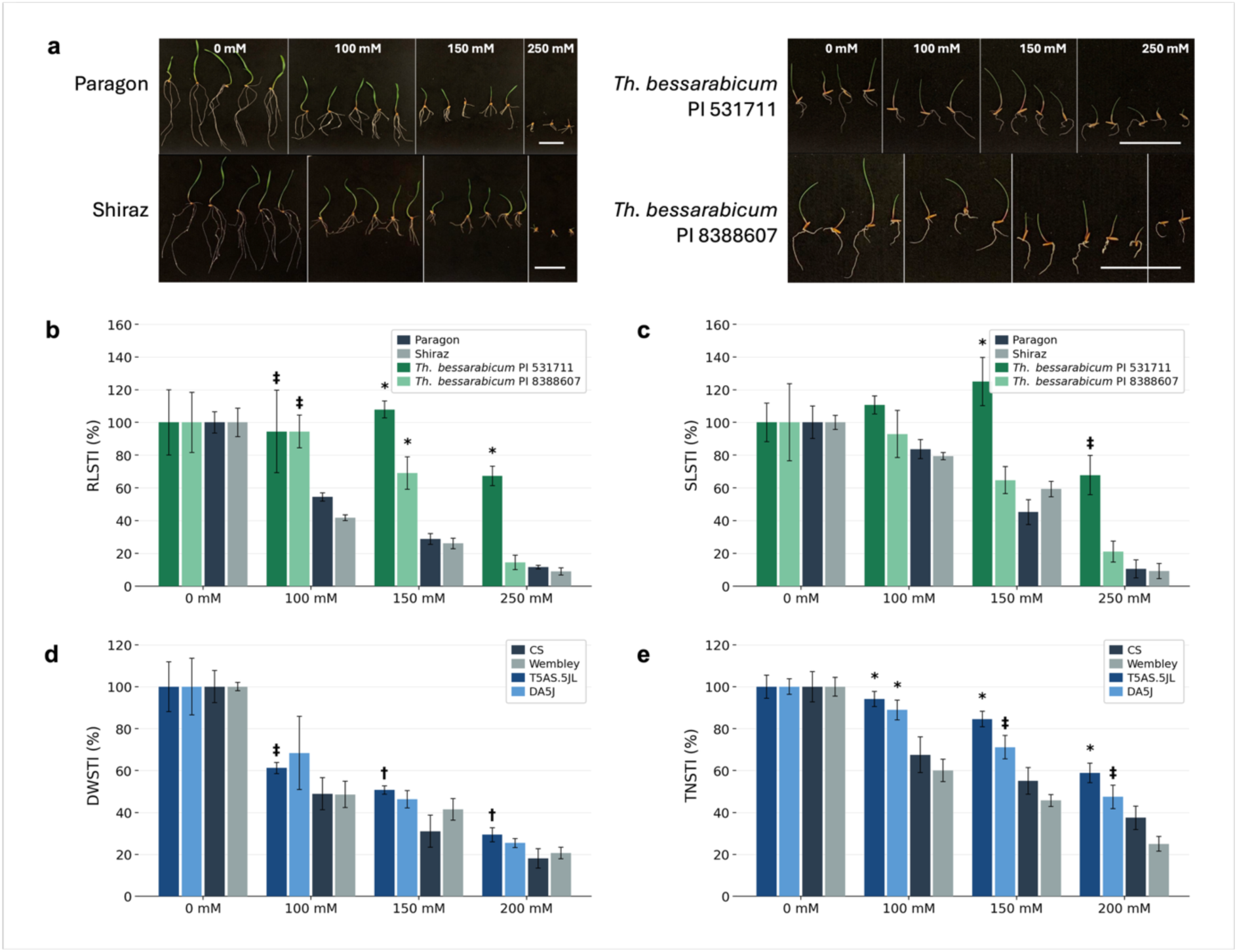
Salt tolerance phenotyping of *Th. bessarabicum* accessions and wheat lines carrying Chr5J introgressions. **(a)** Representative seedlings of Paragon, Shiraz, *Th. bessarabicum* PI 531711 and *Th. bessarabicum* PI 8388607 germinated on filter paper under 0, 100, 150 and 250 mM NaCl for 7 days. Scale bars = 5 cm. **(b)** Root length salt tolerance index (RLSTI) and **(c)** shoot length salt tolerance index (SLSTI) of the same four genotypes at each NaCl concentration. STI expresses each trait measurement as a percentage of the genotype-specific mean under control conditions (0 mM NaCl), normalising for inherent differences in germination vigour between species. **(d)** Shoot dry weight STI (DWSTI) and **(e)** tiller number STI (TNSTI) of bread wheat lines carrying *Th. bessarabicum* Chr5J introgression segments grown in supported hydroponics for 8 weeks under 0, 100, 150 and 200 mM NaCl. T5AS.5JL carries the long arm of Chr5J in a Chinese Spring (CS) background; DA5J carries Chr5J as a disomic addition, also in a CS background. CS and Wembley are salt-sensitive bread wheat controls. All values are means +/- s.e.m. (n = 2-5 per genotype per concentration). * p < 0.05 vs both wheat controls; † p < 0.05 vs CS; ‡ p < 0.05 vs Shiraz (panels b, c) or Wembley (panels d, e); one-sided Mann-Whitney U test.

At the vegetative stage, lines carrying the Chr5J long arm segment showed consistent tolerance benefits in a CS background (**Fig. 6d-e**). Shoot dry weight and tiller number salt tolerance indices (DWSTI and TNSTI respectively) were used to assess biomass and tillering maintenance under progressive NaCl stress. T5AS.5JL, a Robertsonian translocation line, derived from PI 8388607, showed significantly greater DWSTI than Wembley at 100 mM NaCl and significantly greater DWSTI than CS at 150 and 200 mM (all p < 0.05; Fig. 6d). For TNSTI, T5AS.5JL significantly outperformed both CS and Wembley at all three stress concentrations tested (100, 150 and 200 mM; p < 0.05; Fig. 6e). The disomic addition line DA5J, also derived from PI 8388607, showed significantly greater TNSTI than both controls at 100 mM (p < 0.05) and significantly greater TNSTI than Wembley at 150 and 200 mM (p < 0.05), though differences vs CS did not reach significance at the higher concentrations. No significant differences in DWSTI were detected for DA5J at any concentration after correction. Together, these results demonstrate that both T5AS.5JL and DA5J confer measurable improvements in tiller number maintenance under NaCl stress relative to bread wheat controls, with T5AS.5JL showing the most consistent effects across concentrations and traits.

## Discussion

The prevailing assumption that *Th. bessarabicum* contributed the J genome to polyploid *Thinopyrum* species has shaped breeding strategy and genomic study of this lineage for decades, rooted in cytological morphology and ambiguous hybridisation data^7^. Our phylogenomic analysis of 210 single-copy orthologous genes, combined with independent *k*-mer-based genome similarity metrics, decisively refutes this hypothesis (**Fig. 2**). *Th. bessarabicum* clusters robustly with *Th. elongatum* as a sister J/E diploid lineage, well separated from the subgenomes of *Th. intermedium*, whose J-designated subgenome shows phylogenetic affinity to *Aegilops*-related lineages. Sun et al. (2025)^19^ excluded *Th. elongatum* as a genomic contributor to *Th. intermedium* but did not include *Th. bessarabicum*, leaving the diploid J genome untested against either hexaploid or decaploid *Thinopyrum* lineages. The assembly presented here closes this gap: IBSpy *k*-mer analysis against the J genome reference shows all polyploid accessions tested, including decaploid *Th. elongatum*, decaploid *Th. ponticum* and hexaploid *Th. intermedium*, display variation scores exceeding 500, consistent with different-species divergence under established IBSpy thresholds^34, 35^. While broader sampling of polyploid *Thinopyrum* germplasm would strengthen this conclusion, these results are fully consistent with our phylogenomic analysis and provide the first direct sequence-level evidence that the diploid J genome did not contribute to either the hexaploid or decaploid polyploid *Thinopyrum* lineages tested. This finding is independently supported by cytological analysis: differences in rDNA loci distribution and hybridisation patterns between *Th. bessarabicum* J chromosomes and J-designated chromosomes of *Th. intermedium* and *Th. ponticum* suggested the diploid was probably not involved in the origin of these polyploids^41^. The J genome designation in polyploid *Thinopyrum*, assigned on morphological criteria, does not reflect evolutionary descent from the diploid J genome. This discrepancy exemplifies a broader challenge in polyploid Triticeae nomenclature, where convergent chromosome morphology and shared repetitive landscapes can mislead cytogenetic subgenome assignment. We therefore propose that the J subgenome designation in *Th. intermedium* and related polyploid *Thinopyrum* species be revisited in light of the genomic evidence now available, following the phylogenomically informed framework applied here and in Sun et al. (2025)^19^. The practical implication is direct: the J subgenome designation cannot be used as a proxy for shared gene content between *Th. bessarabicum* and polyploid *Thinopyrum* species, and genomic resources developed for one cannot be assumed transferable to the other.

The reciprocal translocation between homoeologous chromosomes 4 and 5 is a defining structural feature of the J genome, with molecular marker evidence previously confirming its presence in *Th. bessarabicum*^30, 42^. We now provide the first gene-level resolution of the breakpoints in a six-species comparative framework (**Fig. 3**). The breakpoint-flanking gene pairs on Chr4J (*WD3L*/*PME*IL1) and Chr5J (*ASA1*/*bHLH*) are identical to those reported in *D. villosum*^36^ and are conserved in *T. urartu*, pointing to a single ancestral rearrangement predating the divergence of the J, V, and A genome lineages. The unrearranged configuration in *Th. elongatum*, *Ae. tauschii*, and *Th. intermedium*, and the additional 4/7 translocation unique to *D. villosum* which post-dates the shared 4/5 event^36^, corroborate this interpretation. Notably, the 4/5 translocation is also present in the wheat A genome^42–44^, reinforcing the inference of a deep common ancestor carrying this rearrangement across J, A, and V lineages while B and D genome progenitors retained the ancestral arrangement. This interpretation is independently supported by Sun et al. (2025)^19^, who identified the same 4/5 rearrangement in the V subgenome of *Th. intermedium* and *Roegneria kamoji*, with breakpoints matching those in *D. villosum*, confirming that the translocation is a shared feature of the V lineage and is absent from the *Th. intermedium* “J” subgenome. The practical significance is illustrated by translocation line T4BS.4BL-4JL, characterised by Patokar et al. (2016)^31^ as carrying a distal 4JL segment, based on homoeologous collinearity with wheat 4BL. Our skim-sequencing analysis instead identifies Chr5J gain, an assignment that is only resolvable with a *Th. bessarabicum* reference genome and an understanding of the 4/5 translocation.

The J genome at 5.87 Gb is approximately 1.33 Gb larger than the sister E genome of *Th. elongatum* (4.54 Gb). Our LTR retrotransposon analysis identifies a recent burst of transposition as the principal driver: 89.6% of intact J genome LTR elements inserted within the last 2 Mya, compared with only 11.2% of intact E genome elements (**Fig. 4c**). However, insertion age estimates derived from LTR divergence are sensitive to the assumed nucleotide substitution rate, which remains uncertain in grasses and may differ between species^45^. Copia elements show genome-wide amplification while Gypsy elements show pronounced centromeric enrichment. The dense accumulation of recently inserted, highly similar Gypsy elements in J genome centromeres provides extensive substrate for recombination between non-homologous centromeres, offering a mechanistic explanation for the elevated frequency of centric fusions and Robertsonian translocations observed in wheat x *Th. bessarabicum* hybrid progeny^30–32^ (**Supplementary Fig. 13**).

*Thinopyrum bessarabicum-*derived wheat germplasm has long demonstrated tolerance to high salinity stress^46, 47^. Amphiploids between *Th. bessarabicum* and wheat survive and set seed at 250 mM NaCl^24, 48^, and chromosomes 2J and 5J were identified early as the primary carriers of salt tolerance loci^23, 26, 49^. Our genome-wide gene content analysis provides a mechanistic basis for that physiological capacity (**Fig. 5**; **Supplementary Table 19**). The most prominent signal is quantitative expansion of broadly conserved gene families shared with bread wheat rather than acquisition of unique salt-specific gene families. Stress transcription factors dominate: NAC family genes show approximately 8-fold enrichment and DREB/AP2/ERF genes show approximately 5-fold enrichment over wheat (both p < 0.001), consistent with independent characterisation of 116 NAC genes with 14 salt-responsive members in the closely related *Th. elongatum*^50^ and the broader pattern of NAC amplification across salt-tolerant *Thinopyrum* species^51^. On the Chr5J long arm, a 52-gene germin-like protein (GLP) tandem array (OG0000191) at approximately 772 Mb represents an 8.84-fold expansion over the bread wheat mean. Synteny analysis shows this locus is collinear with wheat chromosome 4B, which carries the largest TaGLP cluster in bread wheat^52^. GLP enzymes contribute to quantitative stress resistance in cereals through H_2_O_2_-mediated cell wall reinforcement^52–55^, and overexpression of GLP from the halophytic grass *Puccinellia tenuiflora* confers salt tolerance in transgenic *Arabidopsis thaliana*^56^. Across the 29-species panel, OG0000191 copy number is substantially higher in wild grasses than domesticated cereals (mean 26 versus 6; **Supplementary Fig. 10c**), suggesting domestication may have inadvertently reduced this stress response component. Together, the Chr5J stress gene hotspot at 758 to 780 Mb, two flanking NAC arrays at approximately 626 Mb and 725 Mb, and co-localised LEA-2 and transporter orthogroups provide a molecular basis for the chromosome 5 salt tolerance signal identified by decades of cytogenetic work^23, 26, 47^. This quantitative, polygenic expansion strategy is consistent with the proposal that halophytic salt tolerance is seldom attributable to single novel genes but rather to regulatory rewiring and dosage amplification of conserved stress systems^57^. In contrast to the marked transcription factor expansions, genes encoding the canonical Na^+^ exclusion and compartmentalisation machinery, including CHX-type cation/H^+^ exchangers and sodium transporters, showed modest copy number enrichment relative to bread wheat (1.35-fold; 86 orthogroups), with the majority of orthogroups carrying equivalent copy numbers in both species. This suggests that the distinguishing feature of *Th. bessarabicum* salt tolerance at the genomic level is not dosage amplification of ion transport effectors but rather elaboration of the transcriptional regulatory networks that control their expression under ionic stress.

The phenotypic data reported here provide direct experimental validation of the Chr5J salt tolerance signal and place it in the context of the *Th. bessarabicum* accessions from which the lines were derived. At germination stage, accession PI 531711 maintained root and shoot growth at or above unstressed control levels up to 150 mM NaCl, with RLSTI and SLSTI values significantly greater than both bread wheat cultivars at 150 and 250 mM NaCl. This strong germination-stage tolerance is consistent with the characterisation of *Th. bessarabicum* as a halophyte capable of withstanding up to 350 mM NaCl through restriction of sodium and chloride accumulation in leaf tissue^22^. The accession used for genome assembly (PI 531712) could not be evaluated directly due to poor germination under controlled conditions, a difficulty commonly encountered with wild perennial accessions, and phenotyping was conducted on two further accessions from the germplasm collection. At the vegetative stage, direct phenotyping of *Th. bessarabicum* accessions was not possible for the same reason, and assessment was conducted using wheat introgression lines carrying defined Chr5J segments. The improved growth maintenance under NaCl stress observed in both T5AS.5JL and DA5J corroborates earlier cytogenetic evidence that chromosome 5J carries dominant salt tolerance gene(s)^23, 26, 47^. The more consistent significance observed for T5AS.5JL across traits and concentrations relative to DA5J likely reflects the fitness advantage of the Robertsonian translocation, which maintains a euploid 42-chromosome complement, over the disomic addition which carries an extra chromosome pair and the associated dosage imbalance across homoeologous group 5 loci. The J genome assembly now localises this to a defined genomic interval on 5JL, providing candidate gene resolution for a chromosomal salt tolerance signal that has been recognised for nearly four decades but could not previously be attributed to specific loci.

The resource limitations of GISH, FISH, and SNP array analysis have been a persistent constraint on deploying *Th. bessarabicum* diversity in wheat^30^. The reference genome addresses this directly by enabling physical mapping of 277 KASP markers at a median inter-marker spacing of 11.6 Mb and a dual-reference skim-sequencing pipeline that resolves introgression boundaries at 1 Mb and corrects multiple cataloguing errors in existing germplasm (**Supplementary Table 21**). The *Th. bessarabicum* reference genome, introgression line resources and marker framework reported here collectively provide the genomic infrastructure needed to translate the salt tolerance and stress resilience of this wild halophyte into tangible gains for wheat improvement in an era of increasing soil salinity and climate instability.

## Materials and Methods

### Plant material and growth conditions

*Thinopyrum bessarabicum* (accessions PI531711 and PI531712), decaploid *Th. elongatum* (accession PI401007) and *Th. ponticum* (accession PI547312) were obtained from the United States Department of Agriculture (USDA) while *Th. bessarabicum* accession PI 8388607 was obtained from the Germplasm Resource Unit (GRU, John Innes Centre). Genebank introgression lines used for skim-sequencing were obtained from the GRU and from CIMMYT (details in **Supplementary Table 21**).

All plants in this study were grown under controlled glasshouse conditions (18-25°C; 16 h light, 8 h dark) in 2 L pots containing John Innes No. 2 compost.

### Genomic DNA extraction and sequencing

For HMW DNA extraction for genome assembly, approximately 2 g of young leaf tissue from a plant of *Th. bessarabicum* accession PI531712 was harvested following 48 h of dark treatment, snap-frozen in liquid nitrogen and ground to a fine powder under cryogenic conditions. HMW DNA was isolated using a modified Qiagen Genomic DNA protocol^58^, previously adapted for wheat wild relatives^59^. SMRTbell genomic libraries were prepared from purified DNA using PacBio SMRTbell Express Template Prep Kit v3.0, size-selected (15-20 kb insert) using a BluePippin system (Sage Science), and sequenced on a PacBio Revio platform using four SMRT cells by Novogene UK, generating approximately 312 Gb (∼54x coverage) of HiFi reads (**Supplementary Figure S1a-b**; **Supplementary Table 1**). A Hi-C library was prepared from the same individual using the Proximo Hi-C kit for plant tissues (Phase Genomics, USA) and sequenced on an Illumina NovaSeq X Plus (2 x 150 bp) to 112x coverage by Novogene UK (**Supplementary Table 2**).

For downstream genetic and genomic applications, leaf material was collected from two-week-old plants, freeze-dried and ground at 25 Hz for 4 minutes using a TissueLyser II (Qiagen). DNA was extracted using the protocol of Yang et al. (2026)^60^. For whole-genome sequencing applications (skim-sequencing and IBSpy) and probe preparation for *in situ* hybridisation analysis, an additional purification step was performed: the aqueous phase was treated with RNase A and extracted with an equal volume of phenol:chloroform:isoamyl alcohol (25:24:1, pH 8.0) followed by ethanol precipitation. DNA quality was assessed by NanoDrop spectrophotometry. Paired-end libraries for skim-sequencing and IBSpy WGS were sequenced on the Illumina NovaSeq 6000 (2 x 150 bp) by Novogene UK.

### Assembly and scaffolding

Genome size was estimated from *k*-mer analysis of HiFi reads using Jellyfish v2.2.10^61^ and GenomeScope v2.0^62^ (k=31; **Supplementary Figure S1c**). HiFi reads were assembled *de novo* with Hifiasm v0.19.5^63^. Hi-C reads were trimmed with Trimmomatic v0.39^64^ and aligned to the contig assembly using the Arima Genomics mapping pipeline (https://github.com/ArimaGenomics/mapping_pipeline). Contigs were scaffolded with YaHS v1.2a.2^65^, and the Hi-C contact map was manually curated in PretextView v0.2.5 following the Rapid Curation pipeline^66^. Seven pseudomolecules were assembled; scaffolds with strong Hi-C contact with a pseudomolecule but unplaceable within it were assigned to that chromosome and tagged as unlocalised (unloc), with their lengths included in the reported chromosome lengths (**Supplementary Table 3**). Remaining unanchored scaffolds are reported separately. Chromosomes were named and oriented by synteny with *Th. elongatum*^20^. Contaminant scaffolds were identified with TIARA v1.0.2^67^ and removed; unanchored scaffolds classified as bacterial, prokaryotic or extranuclear were excluded from the final assembly. Organelle genomes were assembled using the Oatk pipeline (https://github.com/c-zhou/oatk).

### Genome annotation

Structural and functional annotation followed the same strategy applied in our previous wheat wild-relative genome projects^59, 68^ using the Robust and Extendable eukaryotic Annotation Toolkit (REAT; https://github.com/EI-CoreBioinformatics/reat) and Minos pipeline (https://github.com/EI-CoreBioinformatics/minos), integrating repeat masking, RNA-Seq and Iso-Seq transcript alignments, cross-species protein alignments and ab initio prediction. Briefly, repetitive elements were identified with the EIRepeat pipeline (https://github.com/EI-CoreBioinformatics/eirepeat). RNA was extracted from six tissue types including 7-day old seedlings (at dawn and dusk), roots, flag leaves and spikes (7 days post-anthesis) and developing grains (15 days post-anthesis). Short-read RNA-Seq libraries were sequenced on Illumina NovaSeq 6000 (**Supplementary Table 6**) and Iso-Seq libraries were sequenced on a PacBio Sequel II (**Supplementary Table 7**). Transcript evidence was aligned with HISAT2 v2.2.1^69^ and minimap2^70^, assembled with StringTie2 v2.1.5^71^ and Scallop v0.10.5^72^ and integrated with Mikado^73^. Cross-species protein evidence from ten Poaceae species (**Supplementary Table 8**) was aligned using Spaln v2.4.7^74^ and Miniprot v0.3^75^. AUGUSTUS^76^ was trained on a high-confidence gene model subset and run with extrinsic hints; predictions were consolidated with EVidenceModeler^77^ and polished with Mikado. Wheat gene models from IWGSC RefSeq v2.1^40^ and pangenome projects^78^ were projected with Liftoff v1.5.1^79^. All gene model sources were merged with Minos into a final non-redundant set (**Supplementary Table 10**). Gene model classification (protein-coding, putative, transposable element; high-confidence HC/low-confidence LC) followed the criteria of Grewal et al. (2024)^59^. Functional annotation was performed using the EIFunAnnot pipeline (https://github.com/EI-CoreBioinformatics/eifunannot), which integrates the AHRD pipeline v3.3.3 with BLASTp^80^ against *Arabidopsis thaliana* TAIR10 and UniProt Viridiplantae protein databases^81^ (e-value ≤1e-5) and InterProScan v5.22.61^82^ domain annotation.

### Assembly and annotation quality assessment

Assembly base-level accuracy was evaluated with Merqury v1.3^83^, using a *k*-mer database built from the HiFi reads with meryl (k=21; **Supplementary Fig. 2a**). Hi-C contact map integrity was visually inspected in PretextView v0.2.5 (https://github.com/sanger-tol/PretextView) following manual curation to confirm pseudomolecule continuity and the absence of misjoins (**Supplementary Fig. 2b**). Assembly and annotation completeness were assessed with BUSCO v5.4.3^84^ against the Poales lineage dataset (n=4,896) in genome and protein modes respectively (**Supplementary Fig. 2c**). Assembly continuity was further evaluated using the LTR Assembly Index (LAI; **Supplementary Fig. 2d**), calculated from the LTR_retriever output described below.

### Repeat and transposable element analysis

Intact LTR-retrotransposons were identified using LTR_retriever v.2.9.0^85^ with LTRharvest and LTR_FINDER_parallel, applied to both the *Th. bessarabicum* J genome and the *Th. elongatum* E genome to enable comparative analysis of transposition dynamics. Insertion ages of intact LTR retrotransposons were estimated from the sequence divergence between paired LTRs using a substitution rate of 1.3 × 10⁻⁸ substitutions per site per year^85, 86^. Subfamily classification used BLAST against ClariTeRep (https://github.com/jdaron/CLARI-TE) and TREP v.2025 databases (≥75% identity, ≥40% query coverage, alignment length ≥500 bp).

To compare the chromosomal distribution of Gypsy and Copia LTR-retrotransposons between the J and E genomes, each chromosome was divided into 100 equal-proportion positional bins. Repeat density per bin was calculated as the summed length of intact LTR elements assigned to that bin expressed as a proportion of bin length, normalising for differences in chromosome length between the J and E genomes. Chromosomal positions were scaled to a 0-100% axis to allow visual comparison across chromosomes of differing absolute lengths.

Centromeric domains were delineated using the density of CEREBA-lineage LTR-retrotransposons, which localise specifically to centromeric regions in barley and related Triticeae^87^. CEREBA density was calculated in non-overlapping 100 kb windows and smoothed using a five-window (500 kb) rolling mean. The centromeric domain for each chromosome was defined as the contiguous region in which smoothed CEREBA density exceeded 30% of the chromosome-specific peak value. This threshold was selected to capture the region of pronounced CEREBA enrichment flanking the density maximum while excluding low-level pericentromeric signal present along chromosome arms and was cross-validated by coincidence with gene-poor regions on all seven chromosomes.

### DNA methylation

5mC methylation was detected natively from PacBio Revio HiFi BAM files. HiFi reads were aligned to the nuclear assembly using pbmm2 (--preset HiFi; MAPQ ≥30; https://github.com/PacificBiosciences/pbmm2), and the aligned BAM was sorted and indexed with SAMtools v1.1.6^88^ using a CSI index to support chromosomes exceeding 512 Mb. Methylation frequency at CpG sites was called using aligned_bam_to_cpg_scores from the pb-cpg-tools package (minimum coverage ≥4x), producing per-site 5mC frequency estimates across the genome.

### Phylogenomic inference and divergence time estimation

Single-copy orthologous groups were identified across 28 Triticeae genomes^19, 20, 34–36, 40, 59, 68, 89–96^ and *Brachypodium distachyon*^97^ using OrthoFinder v2.5^98^, treating each subgenome of polyploid species as an independent taxon. Amino acid alignments were generated via the OrthoFinder MSO workflow, back-translated to codon alignments, and trimmed with Gblocks v0.91b^99^ (codon-specific parameters). Trimmed alignments present in all taxa and exceeding 150 bp were concatenated into a supermatrix of 39 taxa. A maximum likelihood tree was inferred with IQ-TREE v2.1.3^100^ under the LG+G4 amino acid substitution model. Individual gene trees were summarised into a coalescent species tree using ASTRAL-III^101^. Divergence times were estimated with MCMCtree^102^ (PAML v4.9) under the independent rates relaxed clock model, with four internal calibrations derived from TimeTree: *Ae. tauschii* vs *T. urartu* (2.1-8.1 Mya), *Triticum* vs *S. cereale* (4.0-11.7 Mya), *H. vulgare* vs *S. cereale* (6.8-18.3 Mya), and the root (*H. vulgare* vs *B. distachyon*; 29.3-35.5 Mya). Bayesian MCMC sampling used a burn-in of 10,000 followed by 1,000,000 samples. The time-calibrated tree was visualised in FigTree v1.4.3.

### *k*-mer-based genome similarity (IBSpy)

Genome similarity between the diploid *Thinopyrum bessarabicum* J genome and polyploid *Thinopyrum* species was assessed using an assembly-free *k*-mer–based approach implemented in IBSpy^34, 35^. Canonical 31-mers were counted from all available assembled genomes, as well as from raw whole-genome sequencing data of decaploid *Th. elongatum* and *Th. ponticum*, sequenced to 13x (385.2 Gb) and 22x (669.8 Gb) coverage, respectively, and from published whole-genome sequencing data for hexaploid *Th. intermedium*^19^ and *Ae. tauschii*^35^, using KMC v3.1.2^103^ with default parameters. IBSpy variation scores were calculated across non-overlapping 50 kb windows using six independent *k*-mer sets, with analyses performed against two reference sequences: the *Th. bessarabicum* genome and the “J” subgenome of *Th. intermedium*. For each *k*-mer set, window-based variation scores were averaged across the reference genome to obtain a single genome-wide distance metric. Interpretation of IBSpy variation scores followed previously established thresholds^34, 35^. An average variation score below 30 indicates a near-identical accession, scores below 250 indicate accessions belonging to the same lineage of the same species, scores below 500 indicate different lineages within the same species, and scores above 500 indicate different species. These thresholds have been shown to robustly distinguish species and lineage relationships within Triticeae using *k*-mer-based comparisons.

### Comparative synteny and translocation analysis

Chromosome-scale synteny between *Th. bessarabicum* and five Triticeae species (*Th. elongatum*, *Th. intermedium* “J” subgenome, *Ae. tauschii*, *T. urartu*, *D. villosum*) was assessed using pairwise BLASTp reciprocal best hits of CDS sequences, with collinear blocks detected using MCScanX^104^. Karyotype ribbon visualisation of the 4/5 translocation used JCVI^105^ orthologues (cscore ≥0.99) across homoeologous chromosome groups 4 and 5 for all six species, rendered with jcvi.graphics.karyotype. Translocation breakpoints were resolved at microsynteny resolution by identifying breakpoint-flanking genes in collinear anchor files for each pairwise comparison using *Th. elongatum* as the ancestral reference; breakpoint gene identities were validated by BLASTp against UniProt/Swiss-Prot (**Supplementary Table 13**).

### Salt tolerance gene analysis

Salt-relevant gene content was analysed across three orthogroup sets: J-genome private, shared among *Thinopyrum* species but absent from bread wheat, and shared between *Th. bessarabicum* and bread wheat. All analyses were restricted to orthogroups with at least one high confidence (HC) protein-coding gene. Salt-relevant genes were identified by searching AHRD functional descriptions against a curated 15-category keyword dictionary spanning NAC, DREB/AP2/ERF, WRKY and MYB transcription factors, ABC transporters, MFS transporters, ion transport genes (HKT, CHX, K⁺ transporters), aquaporins, peroxidases, ROS scavenging enzymes, germin-like proteins, LEA/dehydrins, V-type ATPase/pyrophosphatases, SWEET transporters and compatible solute biosynthesis genes. The 15 functional categories were defined based on established roles in plant salt tolerance mechanisms. Ion transport categories (HKT, NHX, SOS pathway, V-type ATPase/pyrophosphatase, CHX, K+ transporters) were selected based on their documented functions in Na+ exclusion, vacuolar compartmentation, and K+/Na+ homeostasis in Triticeae and related halophytes^51, 106^. Transcription factor categories (NAC, DREB/AP2/ERF, WRKY, MYB) were selected based on their characterised roles in salt-responsive gene regulation across grasses, including in *Thinopyrum* species where NAC, MYB, AP2/ERF and WRKY families show differential expression under salinity stress^51, 107^. Germin-like proteins were included given their established expansion in halophytes and documented roles in ROS scavenging and cell wall reinforcement under salt stress^55, 56^. Remaining categories (aquaporins, ABC and MFS transporters, peroxidases, LEA/dehydrins, SWEET transporters, compatible solute biosynthesis) were selected on the basis of their broader functional evidence in osmotic adjustment, ROS detoxification, and stress protection across plants under salinity^106^. TE-associated genes were excluded prior to matching. Per-category copy number enrichment in shared orthogroups was tested using the Wilcoxon signed-rank test (one-sided, Benjamini-Hochberg correction) comparing *Th. bessarabicum* HC gene counts to mean per bread wheat subgenome (A, B, D). Genomic distribution was visualised as dual-track heatmaps across 10 Mb chromosome bins.

### Generation of wheat-*Th. bessarabicum* introgression lines

Introgression lines were generated using the crossing strategy described in Grewal et al. (2018)^30^. Hexaploid wheat cv. Paragon *ph1/ph1* was crossed as female with *Th. bessarabicum* PI531712 to produce F1 hybrids, which were recurrently backcrossed to Paragon *Ph1/Ph1*. Plants at BC_1_-BC_3_ generation were genotyped with the Axiom^®^ Wheat-Relative Genotyping Array^108^ to identify those carrying introgressions. Selected heterozygous introgression lines were self-fertilised to homozygosity, with KASP marker-assisted selection replacing the array from BC_4_ generation onwards, enabling co-dominant dosage scoring to distinguish heterozygous from homozygous introgressions.

### KASP marker development and physical localisation

Chromosome-specific KASP markers polymorphic between *Th. bessarabicum* and hexaploid wheat were developed as described in Grewal et al. (2020)^37^. A further 107 markers developed for other wild relative species^38, 39, 109^ were tested against *Th. bessarabicum* (accessions PI531711 and PI531712) in this study. All 277 markers validated as polymorphic were physically localised by aligning SNP-containing probe sequences to the J genome assembly using BLASTn (e-value ≤1e-5), retaining single best-scoring alignments. Physical positions and wheat RefSeq v2.1 coordinates are reported in (**Supplementary Table 20**).

### Genomic *in situ* hybridisation

Metaphase chromosome spreads and multicolour GISH were carried out as described in Grewal et al. (2017)^110^. Genomic DNA from *T. urartu* (A genome) was labelled with Alexa Fluor 488 (green); *Ae. speltoides* (B genome) DNA was fragmented (200-1000 bp) and used as blocking DNA (blue–purple); *Ae. tauschii* (D genome) DNA was labelled with Alexa Fluor 594 (red); and *Th. bessarabicum* DNA was labelled with Alexa Fluor 546 (yellow). Slides were counterstained with DAPI and analysed on a Zeiss Axio ImagerZ2 epifluorescence microscope with Metafer4/ISIS software (Metasystems).

### Skim-sequencing and introgression mapping

Genomic DNA was extracted and sequenced as described above by Novogene UK to 0.05x coverage per sample. Reads were trimmed with Trimmomatic v0.39^64^ and aligned to a concatenated reference comprising bread wheat RefSeq v2.1 and the *Th. bessarabicum* assembly using HISAT2 v2.2.1 (--no-spliced-alignment --no-unal), as described in King et al. (2026)^111^. Properly paired unique alignments (MAPQ ≥10) were retained, duplicates removed with Picard (Broad Institute), and BAM files indexed with SAMtools. Coverage was quantified across non-overlapping 1 Mb windows with BEDTools^112^ and normalised coverage deviation calculated using a custom Python script (available at https://github.com/Surbhigrewal/Introgression_mapping). Introgression intervals were classified as windows showing sustained deviation below 0.4 across ≥10 consecutive wheat bins accompanied by deviation above 0.5 across ≥3 consecutive *Th. bessarabicum* bins.

### Salt tolerance phenotyping

#### Germination-stage screening

Seeds of three *Th. bessarabicum* accessions (PI 531711, PI 531712 and PI 8388607) and two *T. aestivum* cultivars (Paragon, Shiraz) were surface-sterilised in 5% (v/v) sodium hypochlorite for 15 minutes, washed three times with deionised water, and placed in moist Petri dishes at 4°C for 2 days to break dormancy. Five seeds per genotype were then transferred to Petri dishes containing 2.5 ml of NaCl solution at 0, 100, 150 or 250 mM. Dishes were sealed with parafilm and incubated in the dark for 2 days followed by 5 days under a 16 h/8 h light/dark regime at 20°C. After 7 days, seedlings were removed and photographed. Shoot length (SL) and root length (RL) were measured by freehand tracing in ImageJ. A salt tolerance index (STI) was calculated for each trait as the individual trait value expressed as a percentage of the genotype-specific mean under control conditions (0 mM NaCl), normalising for inherent growth rate differences between species.

#### Vegetative-stage hydroponics

Salt tolerance at the vegetative stage was evaluated for T5AS.5JL and DA5J, alongside CS and Wembley as bread wheat controls, in a supported hydroponic system. Seeds were surface-sterilised as above and stratified at 4°C for 5 days to synchronise germination, then transferred to mesh pots filled with Hydroleca clay pellets in randomised position within hydroponic tanks. The nutrient solution was quarter-strength modified Hoagland solution (pH 6.0)^113^, aerated continuously and replaced every 14 days. Growth conditions were 21°C day/15°C night, 16 h photoperiod, 400 µmol m^-2^ s^-1^ PAR. NaCl was applied incrementally from day 2 post-transplant in 25 mM steps twice daily using a 5 M NaCl stock solution until target concentrations of 0, 100, 150 or 200 mM were reached; CaCl_2_ was co-applied to maintain a Na^+^:Ca^2+^ molar ratio of 15:1. Whole shoot biomass was harvested 28 days after full salt treatment, oven-dried at 60°C for 7 days and weighed to determine shoot dry weight (DW). Tiller number (TN) was recorded at the same timepoint. DWSTI and TNSTI were calculated using the same formula as the germination STI. Statistical comparisons were made using one-sided Mann-Whitney U tests against each wheat control. The Chr5J-carrying lines evaluated here, T5AS.5JL and DA5J, both derive from accession PI 8388607 (Supplementary Table 21). CIMMYT-derived lines carrying Chr5J segments were excluded from salt tolerance phenotyping as these are maintained in a Prinia background, which has shown early evidence of salt tolerance and would confound attribution of any phenotypic effect to the introgressed J-chromosome segment.

### Data availability

The raw sequence files for the HiFi, Hi-C, RNA-Seq and IsoSeq reads were deposited in the European Nucleotide Archive (ENA) under study accession PRJEB106499. The raw sequence reads for accessions of decaploid species *Th. elongatum* and *Th. ponticum* are deposited in the ENA under study accession PRJEB112780. The final chromosome-scale assembly consisting of the nuclear and organelle genomes was deposited at ENA. The final assembly, gene models and repeat annotations are also available on figshare. The genome browser is available at GrainGenes^114^.

## Acknowledgements

This work was supported by the Biotechnology and Biological Sciences Research Council (BBSRC) [BB/P016855/1 and BB/X011003/1] as part of the Designing Future Wheat (DFW) and Delivering Sustainable Wheat (DSW) programmes. Studentships were supported by the BBSRC Doctoral Training Programme [BB/T008369/1] for Nikki Walter and the Nottingham Future Food Beacon for Jack Walker. We are grateful for access to the University of Nottingham’s Ada high performance computing (HPC) service. Part of this work was also delivered via Transformative Genomics the BBSRC funded National Bioscience Research Infrastructure (BBS/E/ER/23NB0006) at Earlham Institute by members of the Genomics Pipelines and Core Bioinformatics Groups. We are also grateful to Eric Yao and Taner Sen for hosting the assembly at Graingenes, supported by the US. Department of Agriculture, Agricultural Research Service, Project No. 2030–21000-056-00D.

## Author contributions

N.W. and J.W. contributed equally to this work. S.G., I.P.K., and J.K. designed the project and acquired funding. N.W., J.W., J.N., J.Q-C., E.C-G., S.G.K., W.G. and S.G performed bioinformatics analysis. G.G.K. and D.Sw. performed genome annotation. J.W., C.Y., D.Sc., S.A., and A.S-G. performed DNA extraction. S.G., J.W., C.Y., D.Sc., S.A., A.S-G., I.P.K., and J.K. contributed to germplasm development. C.Y. and K.H. performed genomic *in situ* hybridisation. J.W., D.W., J.A., M.H. and S.G. acquired funding for and planned the phenotyping experiments. J.W carried out the phenotypic analysis. N.W., J.W., and S.G. wrote the paper with input from all authors. S.G. supervised the project.

## Additional information

Supplementary Information includes Supplementary Tables 1-21 and Supplementary Figures 1-13.

**Correspondence** and requests for materials should be addressed to Surbhi Grewal.

