## Supplementary Figures for "Chromosome-scale genome assembly of diploid halophyte *Thinopyrum bessarabicum* excludes J genome from polyploid *Thinopyrum* ancestry"

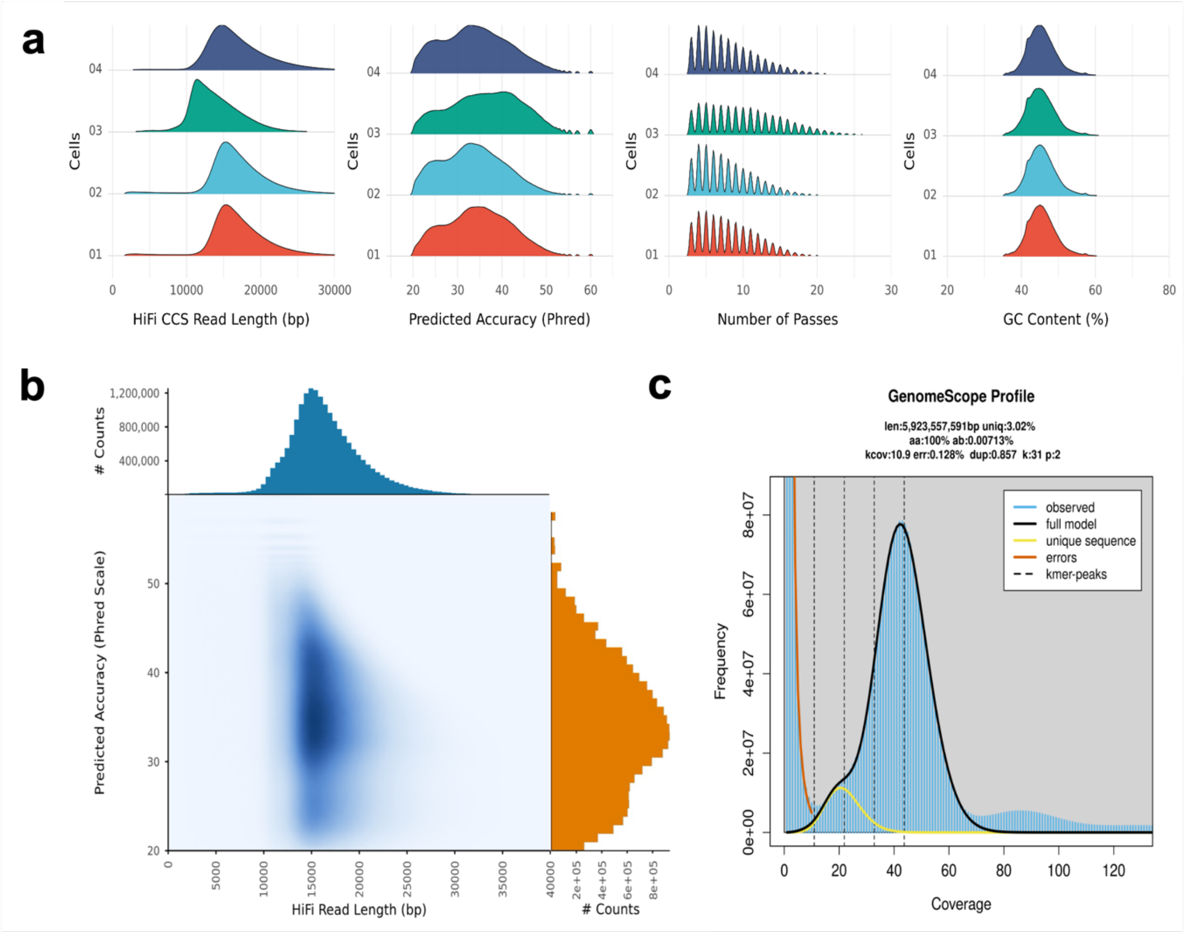


**Supplementary Figure 1. Summary of HiFi reads and genome survey for the *Th. bessarabicum* J genome.**

**(a)** Distribution of HiFi read length, predicted accuracy, number of passes and GC content for each of the 4 PacBio Revio cells. **(b)** Distribution of HiFi read length and predicted accuracy. **(c)** Genome size estimate using GenomeScope2 (k=31).

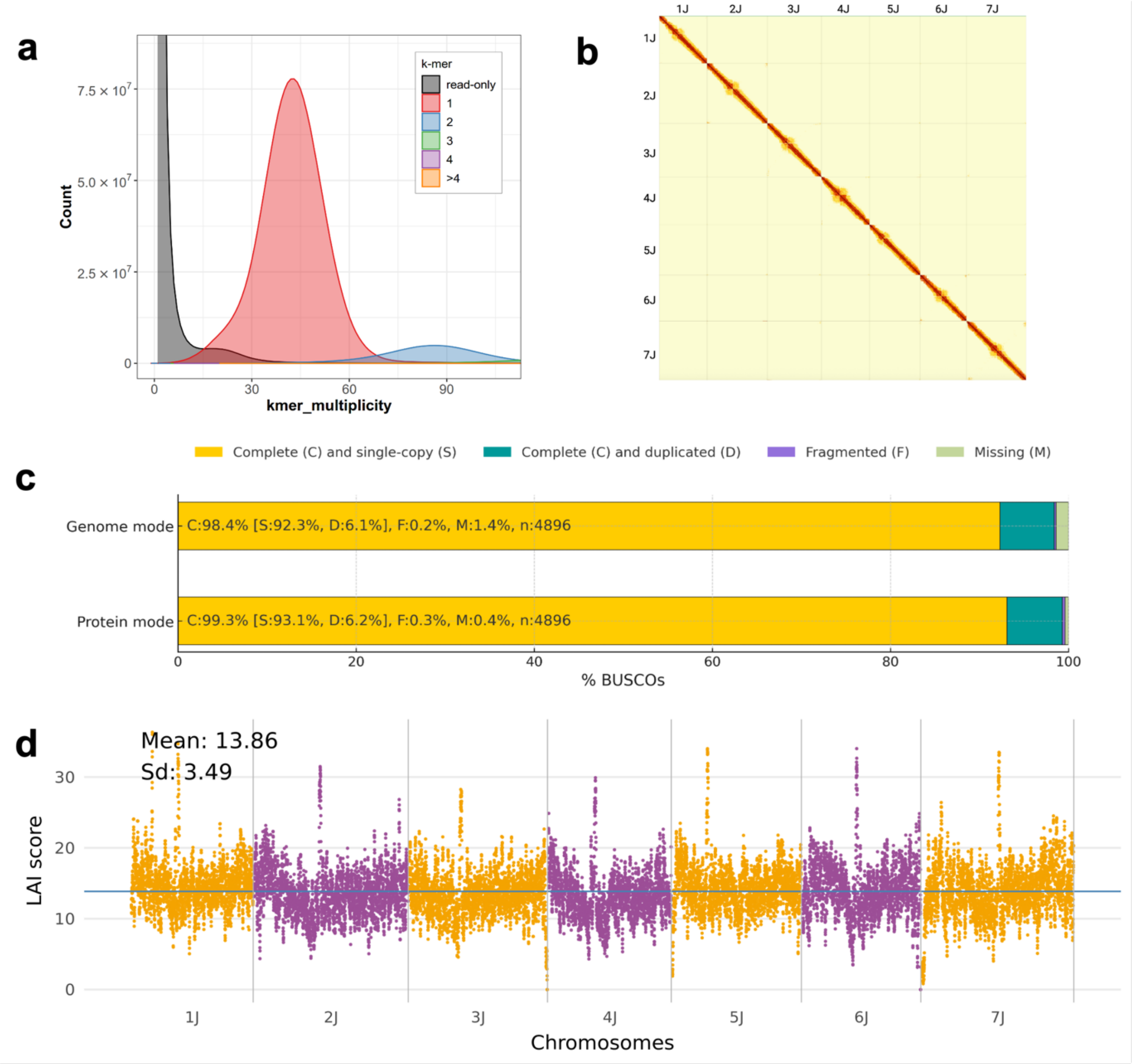


**Supplementary Figure 2. Validation of *Th. bessarabicum* J genome assembly.**
**(a)** Merqury copy number spectrum of the *k*-mers plotted as stacked histograms coloured by the copy numbers found in the primary assembly. The read set *k*-mers absent from the assembly (likely to be sequencing errors in the reads) are plotted in grey, while the assembly *k*-mers absent from the read set (likely to be base errors in the assembly) are plotted as a bar at zero multiplicity, coloured by the copy numbers found in the assembly. **(b)** Chromosome Hi-C contact maps across the 7 pseudomolecules of the J assembly, where the dashed lines indicate pseudomolecule boundaries. **(c)** Completeness of the J assembly as assessed by BUSCO (Simão et al., 2015) using the Poales dataset with a total of 4896 groups. **(d)** Evaluation of J genome assembly by LTR Assembly Index (LAI) scores which were calculated using 3 Mb-sliding windows with 300-kb steps and plotted as dots. The blue line indicates average LAI score for the whole genome.

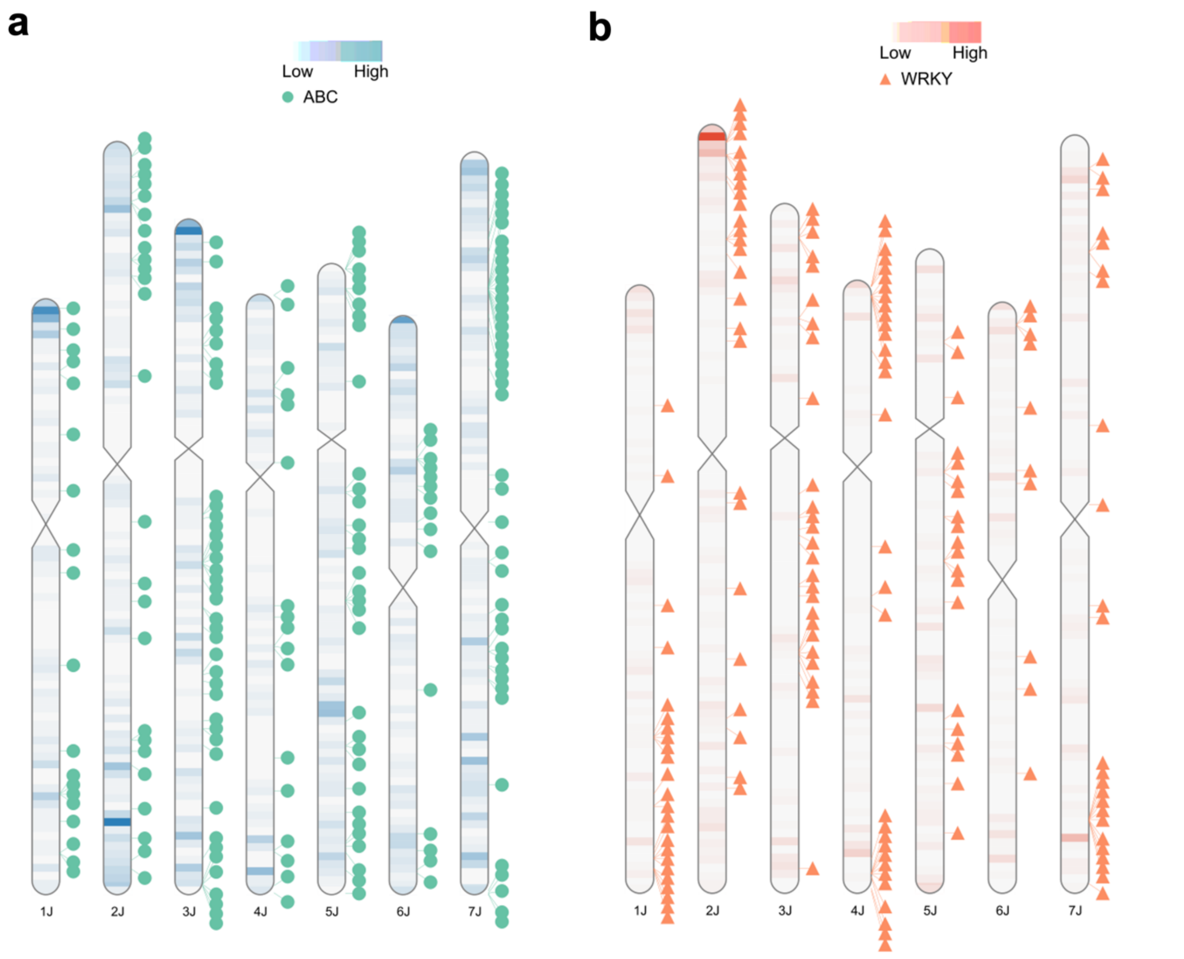


**Supplementary Figure 3. Distributions of disease resistance-related genes across the whole *Th. bessarabicum* J genome.**
**(a)** The overlaid heatmap shows the density of defence-related kinase domain containing genes (Supplementary Table 12), and the circle tack labels refer to distribution of ABC transporter genes. **(b)** The overlaid heatmap shows the density of Cytochrome P450 genes (Supplementary Table 12), and the triangle tack labels refer to distribution of WRKY DNA-binding domain containing genes. Different gene categories extract using InterPro IDs.

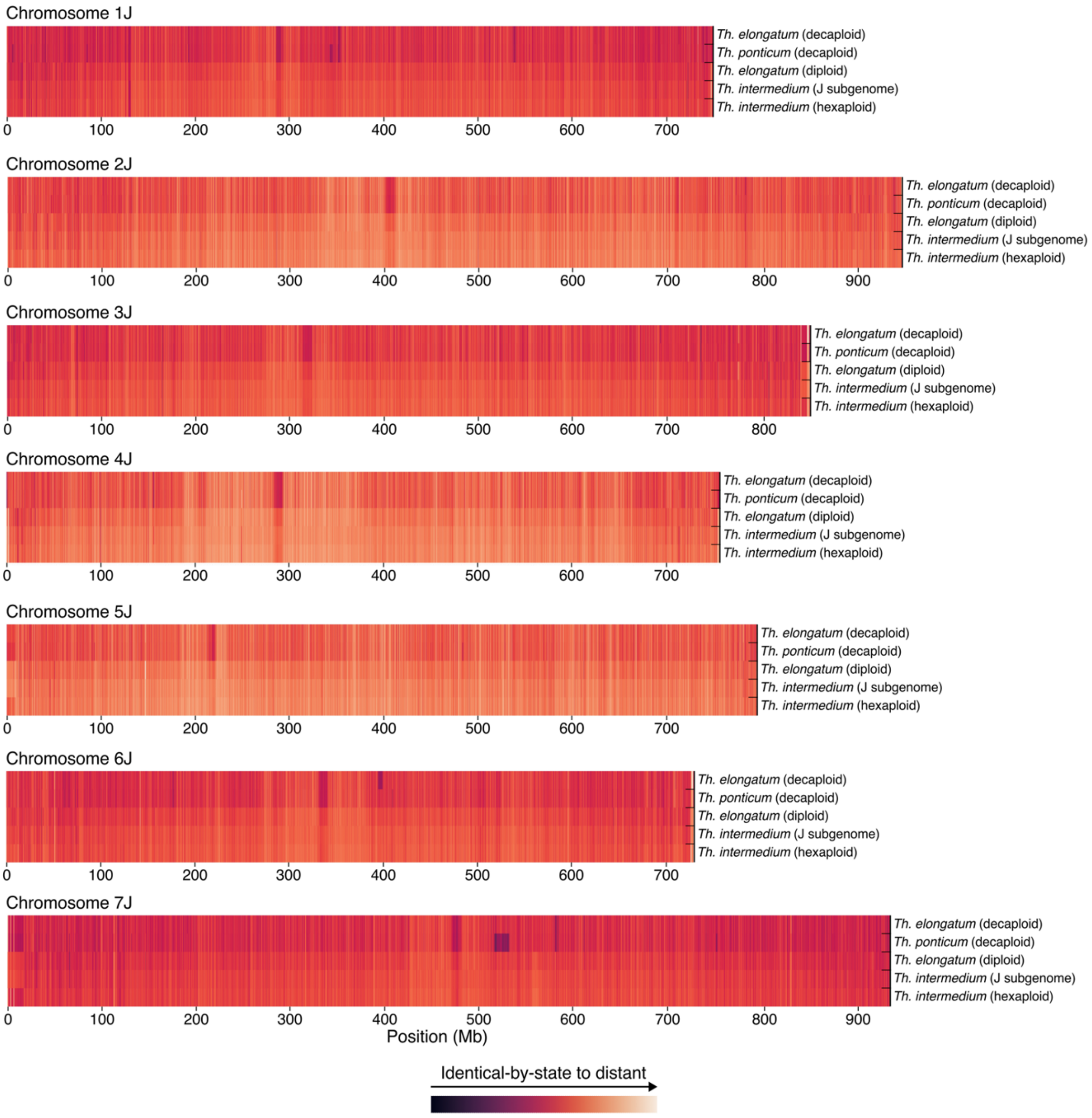


**Supplementary Figure 4. Chromosome-resolved *k*-mer similarity profiles across the *Th. bessarabicum* genome.**

Heatmaps showing chromosome-wide *k*-mer similarity between the *Th. bessarabicum* genome (reference) and selected Thinopyrum accessions. Each panel corresponds to one chromosome (1J–7J), with genomic position shown along the x-axis (Mb). Rows represent comparisons with *Th. elongatum* (decaploid), *Th. ponticum* (decaploid), diploid *Th. elongatum*, *Th. intermedium* (J subgenome), and hexaploid *Th. intermedium*. Colours represent IBSpy similarity scores ranging from identical-by-state (dark) to increasingly divergent *k*-mer profiles (light), as indicated by the colour scale.

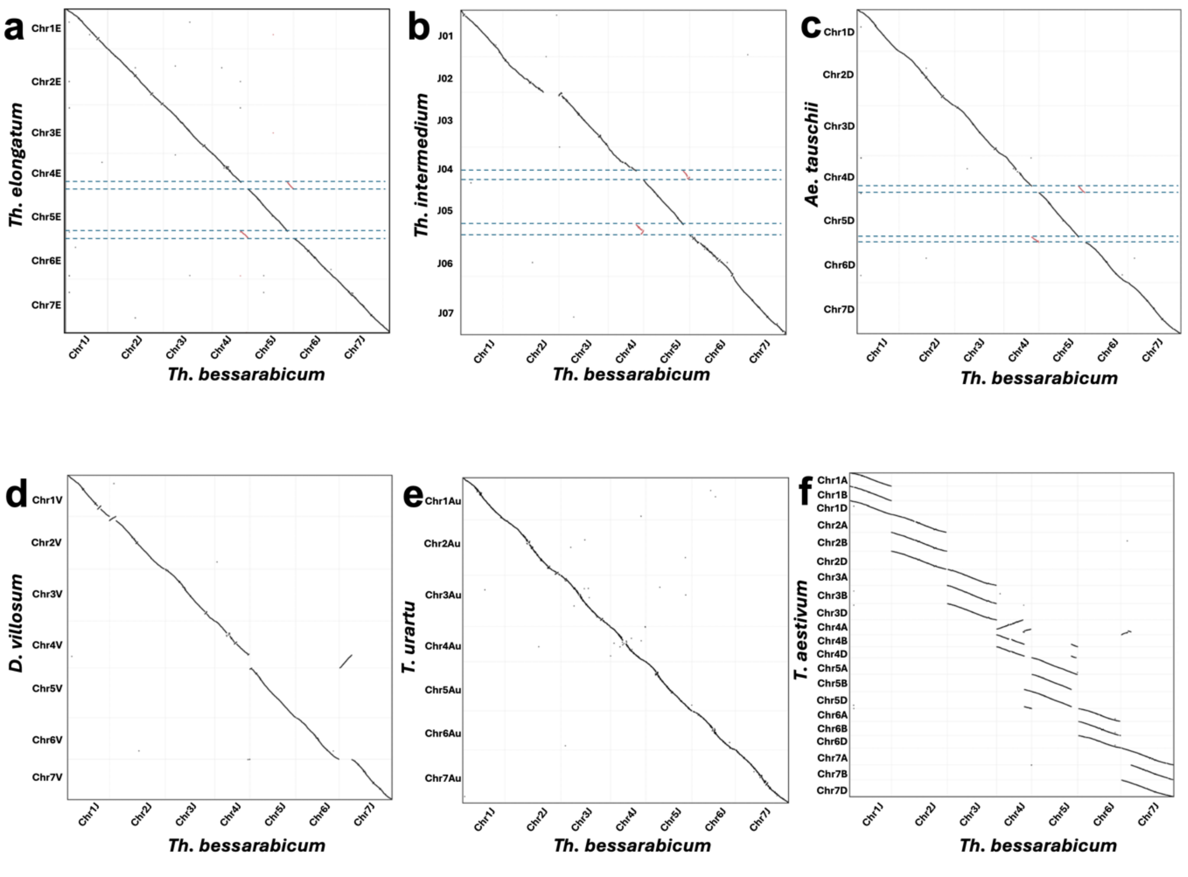


**Supplementary Figure 5. Gene-based sequence alignment comparison between *Th. bessarabicum* J genome** and **(a)** *Thinopyrum elongatum* (E); **(b)** *Th. intermedium* ("J" subgenome); **(c)** *Ae. tauschii* (D) Aet_v4.0; **(d)** *D. villosum* (V); **(e)** *T. urartu* ($A^u$); **(f)** *T. aestivum* RefSeq2.1. Each dot represents a collinear gene pair between the two genomes. Black dots indicate collinear gene pairs on the same homoeologous chromosome. Red dots indicate gene pairs within the chromosomal segment translocated between chromosomes 4J and 5J in *Th. bessarabicum*; in panels A (*Th. elongatum*), B (*Th. intermedium* J subgenome), and C (*Ae. tauschii*), these genes align to their ancestral chromosome positions, confirming that these species retain the pre-translocation arrangement. Black dashed lines mark the translocation breakpoint positions on the *Th. bessarabicum* chromosomes. Panels D–F show species sharing the 4/5 translocation with *Th. bessarabicum*; synteny is continuous across the breakpoints and no dashed lines are shown. In panel D (*D. villosum*), an additional off-diagonal signal between Chr4V and Chr7V reflects the V-genome-specific 4/7 translocation, absent in both the J and E genomes. In panel F (*T. aestivum*), the fragmented multi-segment pattern reflects the complex rearrangement history of the wheat A subgenome, including a 4AL/5AL translocation inherited from *T. urartu* and subsequent lineage-specific rearrangements including a pericentric inversion of 4A and a 4/7 translocation.

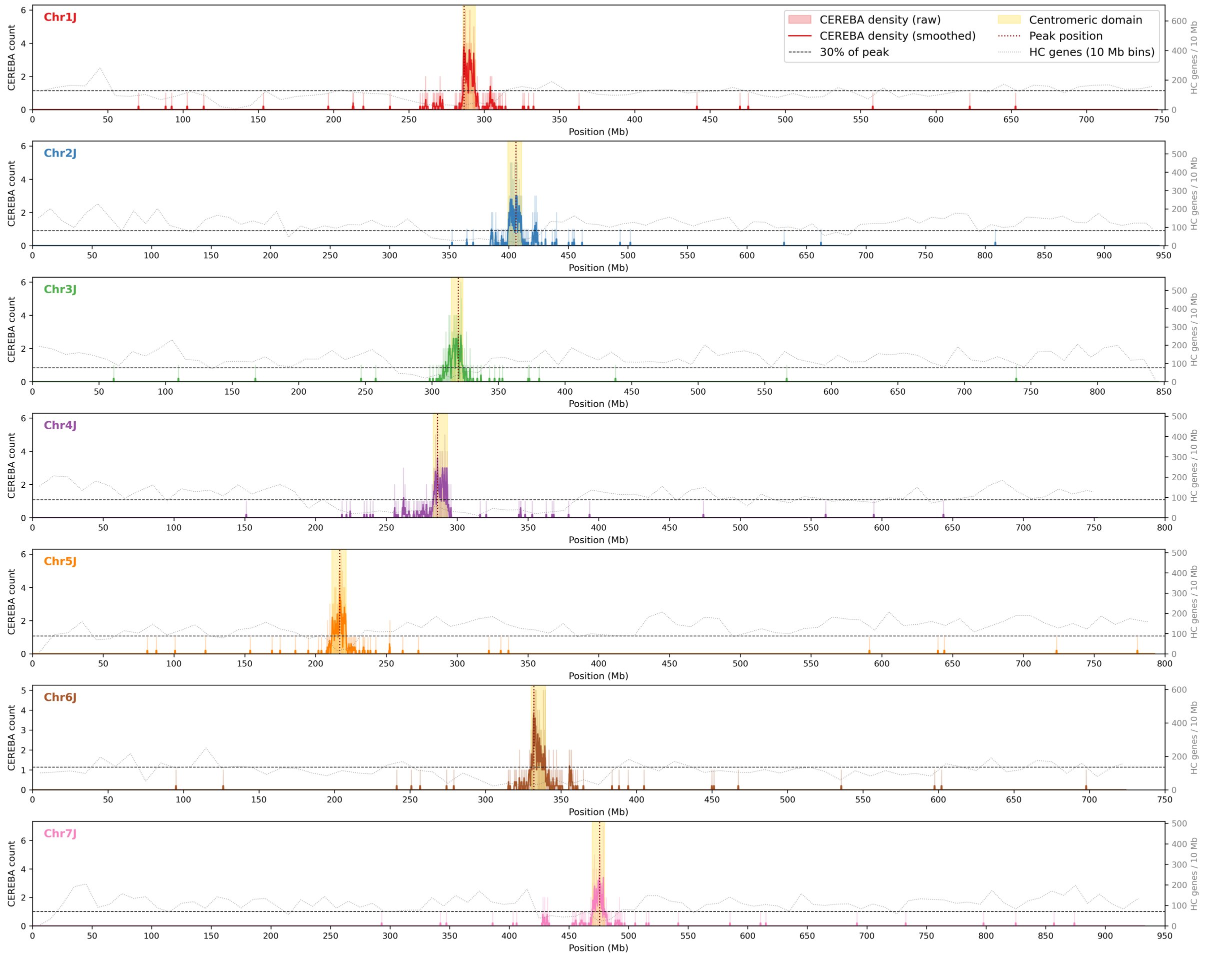


**Supplementary Figure 6. CEREBA-based centromeric domain profiles across the seven *Th. bessarabicum* J genome chromosomes.**

CEREBA-lineage LTR-retrotransposon density was calculated in non-overlapping 100 kb windows along each pseudomolecule. Raw counts are shown as filled bars (chromosome-specific colour); the smoothed signal (500 kb rolling mean across five consecutive windows) is shown as a solid line in a darker shade of the chromosome colour, overlying the raw density bars. The horizontal dashed line marks the 30% of peak density threshold used for centromeric boundary calling. Gold-shaded regions indicate the inferred centromeric domain for each chromosome; the vertical dotted line marks the CEREBA density peak position. High-confidence (HC) protein-coding gene density in 10 Mb bins is overlaid on the secondary y-axis (grey dashed line). Centromeric coordinates and CEREBA element counts are given in Supplementary Table 16. The legend is shown for Chr1J and applies to all panels.

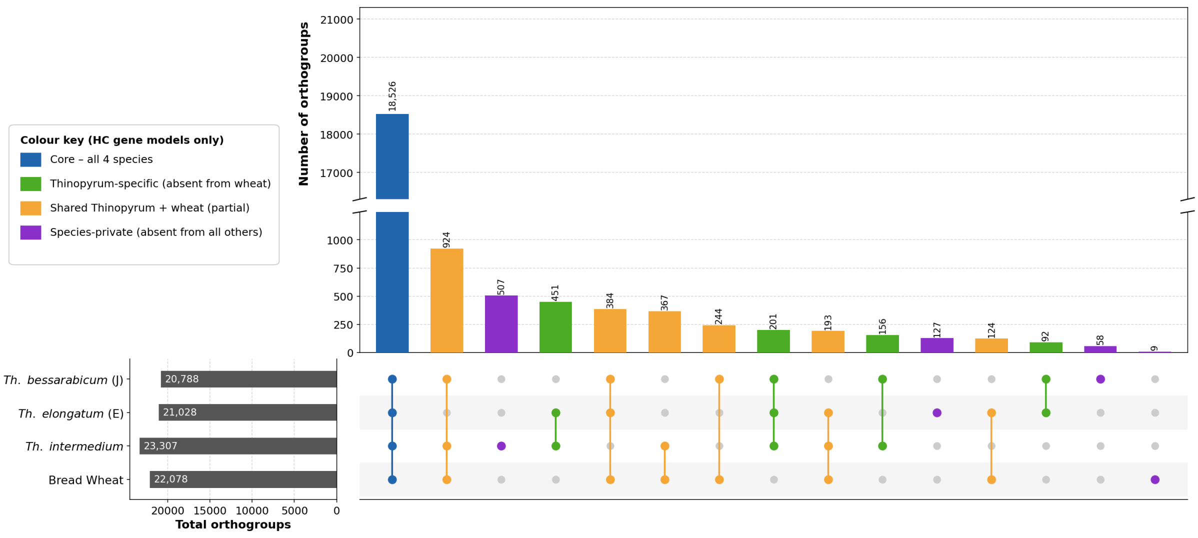


**Supplementary Figure 7. Orthogroup membership across *Th. bessarabicum*, *Th. elongatum*, *Th. intermedium*, and bread wheat.**

UpSet plot showing the number of high-confidence (HC) orthogroups in each intersection of the four focal species. Bar height indicates the number of orthogroups in each intersection; horizontal bars on the left show the total number of orthogroups per species. Orthogroup membership is defined by presence of at least one HC gene model in that species, with *Th. intermedium* and bread wheat treated as single units by collapsing their contributing subgenomes (J, St, and V for *Th. intermedium*; A, B, and D for bread wheat). Species-private orthogroups (absent from all other species in the full 38-species panel) are shown in purple; orthogroups shared across all three *Thinopyrum* species but absent from bread wheat are shown in green; orthogroups shared between *Thinopyrum* and bread wheat are shown in orange; and orthogroups shared across all four focal species are shown in blue. The three gene universes analysed for salt-relevant content (Supplementary Tables 17-19) correspond to the species-private, wheat-absent, and core shared intersections respectively.

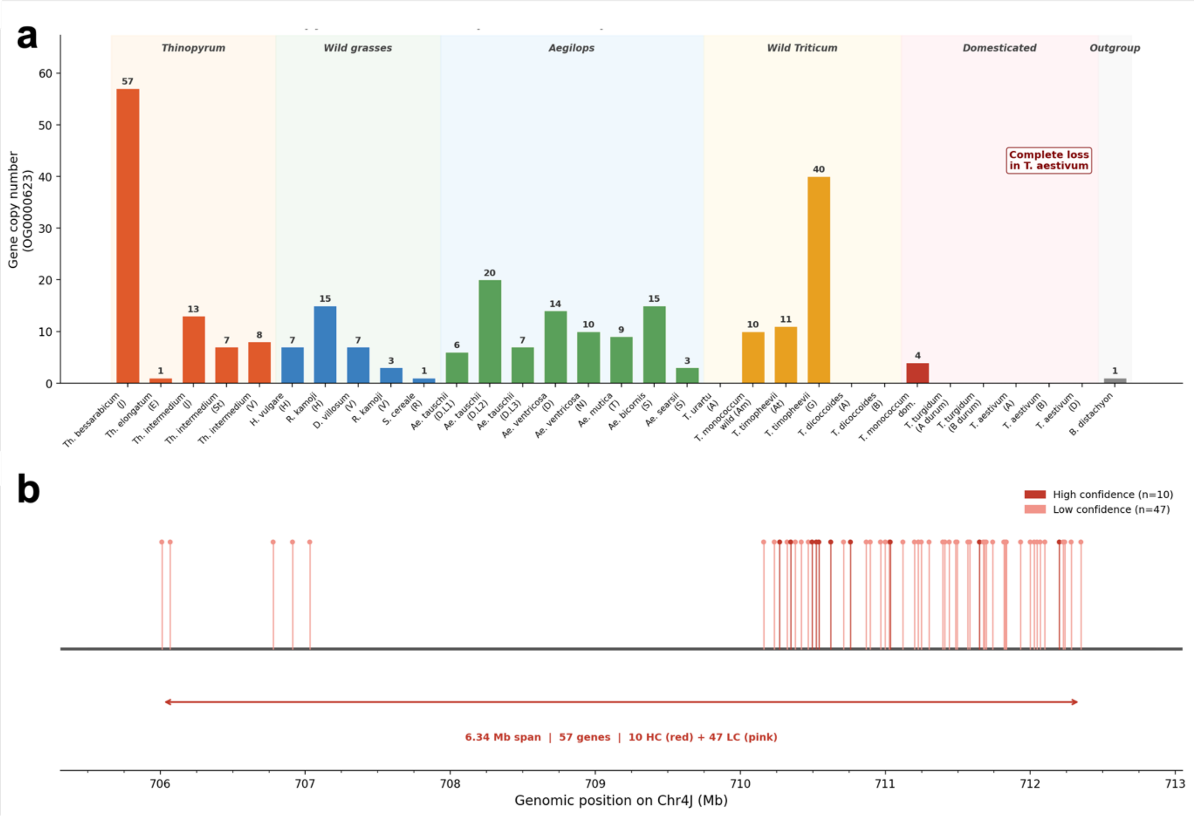


**Supplementary Figure 8. OG0000623 NAC domain transcription factor copy number and chromosomal organisation across the Triticeae.**

**(a)** Gene copy number for OG0000623 across 38 Triticeae species ordered by phylogenetic group. Background shading indicates species groupings: *Thinopyrum* (orange), wild grasses (green), *Aegilops* (blue), wild *Triticum* (yellow), domesticated cereals (red), and outgroup (grey). *Th. bessarabicum* carries the highest copy number in the panel (n = 57). All three *T. aestivum* subgenomes carry zero copies, indicating complete loss during polyploidisation or domestication. **(b)** Genomic organisation of the OG0000623 tandem array on Chr4J in *Th. bessarabicum*. Each vertical line represents one gene model; high-confidence gene models are shown in dark red (n = 10) and low-confidence models in pink (n = 47). The array spans 6.34 Mb between 706 and 712 Mb.

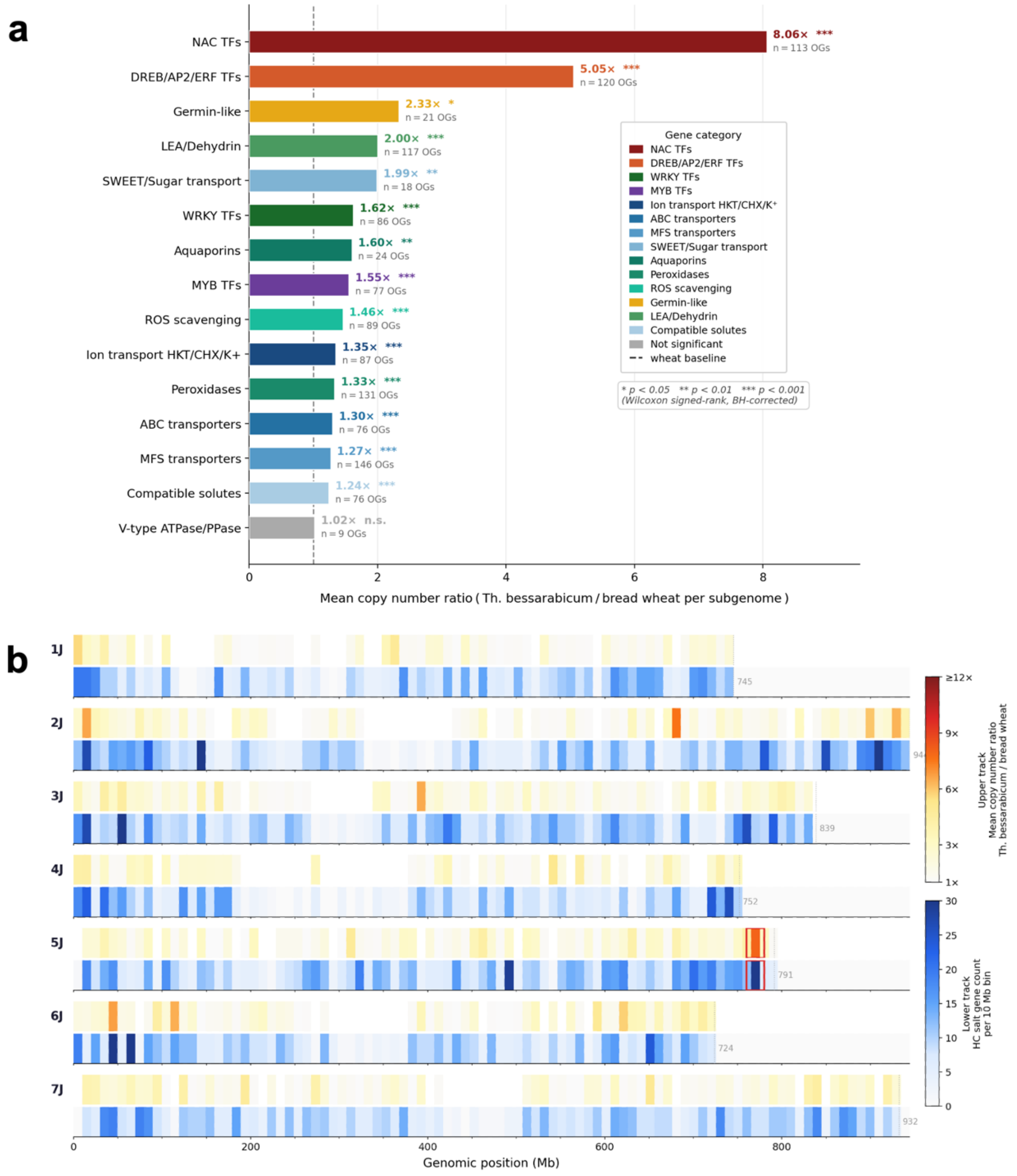


**Supplementary Figure 9. Salt-tolerance gene family enrichment in *Th. bessarabicum* relative to bread wheat.**

**(a)** Mean copy number ratio for 15 salt-relevant gene family categories in *Th. bessarabicum* relative to each bread wheat subgenome, for high-confidence orthogroups shared between the two species (n = 20,078 orthogroups). Bars represent the mean *Th. bessarabicum*/bread wheat copy number ratio per orthogroup within each category. Fourteen of fifteen categories showed significantly higher copy numbers in *Th. bessarabicum* (Wilcoxon signed-rank test, Benjamini-Hochberg correction; significance thresholds indicated by asterisks). The number of orthogroups contributing to each category is shown to the right of each bar. The dashed vertical line indicates the bread wheat baseline (ratio = 1). **(b)** Genome-wide distribution of salt-tolerance gene expansion across the seven *Th. bessarabicum* J-genome chromosomes. Each chromosome is represented by two tracks: the upper track shows the mean *Th. bessarabicum*/bread wheat copy number ratio per 10 Mb bin (warm colours, capped at 12-fold); the lower track shows the count of high-confidence salt-relevant genes per 10 Mb bin (blue scale). Only bins containing at least five genes are shown in the ratio track. The dotted vertical line marks the assembly end of each chromosome; the grey number indicates assembly length in Mb. The red box on Chr5J marks the 760 to 780 Mb region examined in detail in Figure 5a-b.

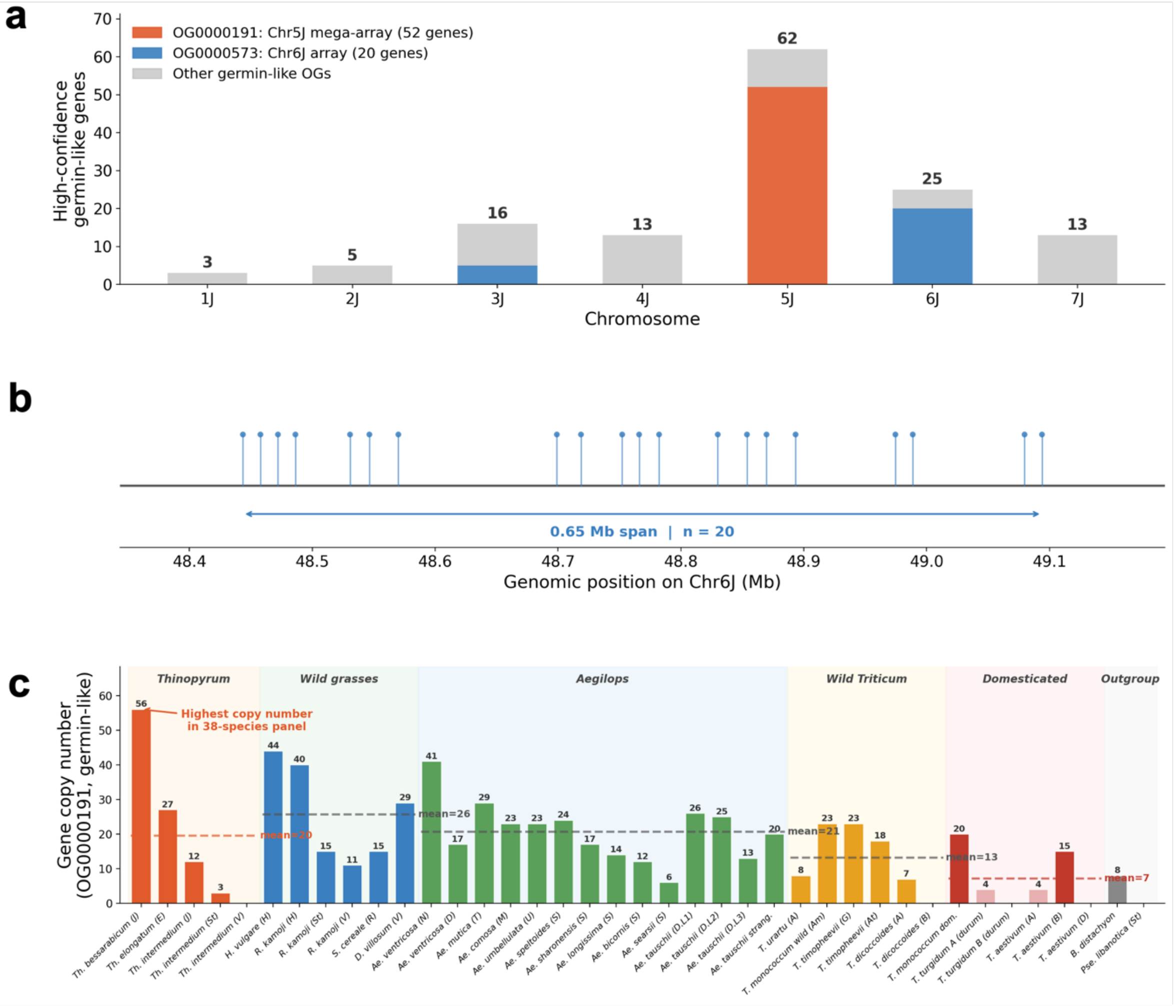


**Supplementary Figure 10. Germin-like protein gene content in *Th. bessarabicum*.**
**(a)** Chromosomal distribution of HC germin-like protein genes across the seven J-genome chromosomes. Bars are coloured by orthogroup: OG0000191 (orange, Chr5J mega-array), OG0000573 (blue, Chr6J array), and all other germin-like orthogroups (grey). Numbers above each bar indicate the total HC gene count per chromosome. **(b)** Genomic organisation of the OG0000573 tandem array on Chr6J. Each vertical line and filled circle represents one HC gene model. The array comprises 20 HC genes spanning 0.65 Mb at approximately 48 to 49 Mb on Chr6J. **(c)** OG0000191 copy number across the 38-species Triticeae panel ordered by phylogenetic group. Background shading indicates species groupings: *Thinopyrum* (orange), wild grasses (green), *Aegilops* (blue), wild *Triticum* (yellow), domesticated cereals (red), and outgroup (grey). Dashed horizontal lines show the group mean copy number. *Th. bessarabicum* carries the highest copy number of any species in the panel (n = 56). The generally lower copy numbers in domesticated cereals relative to wild grasses reflect copy number contraction associated with polyploidisation or domestication.

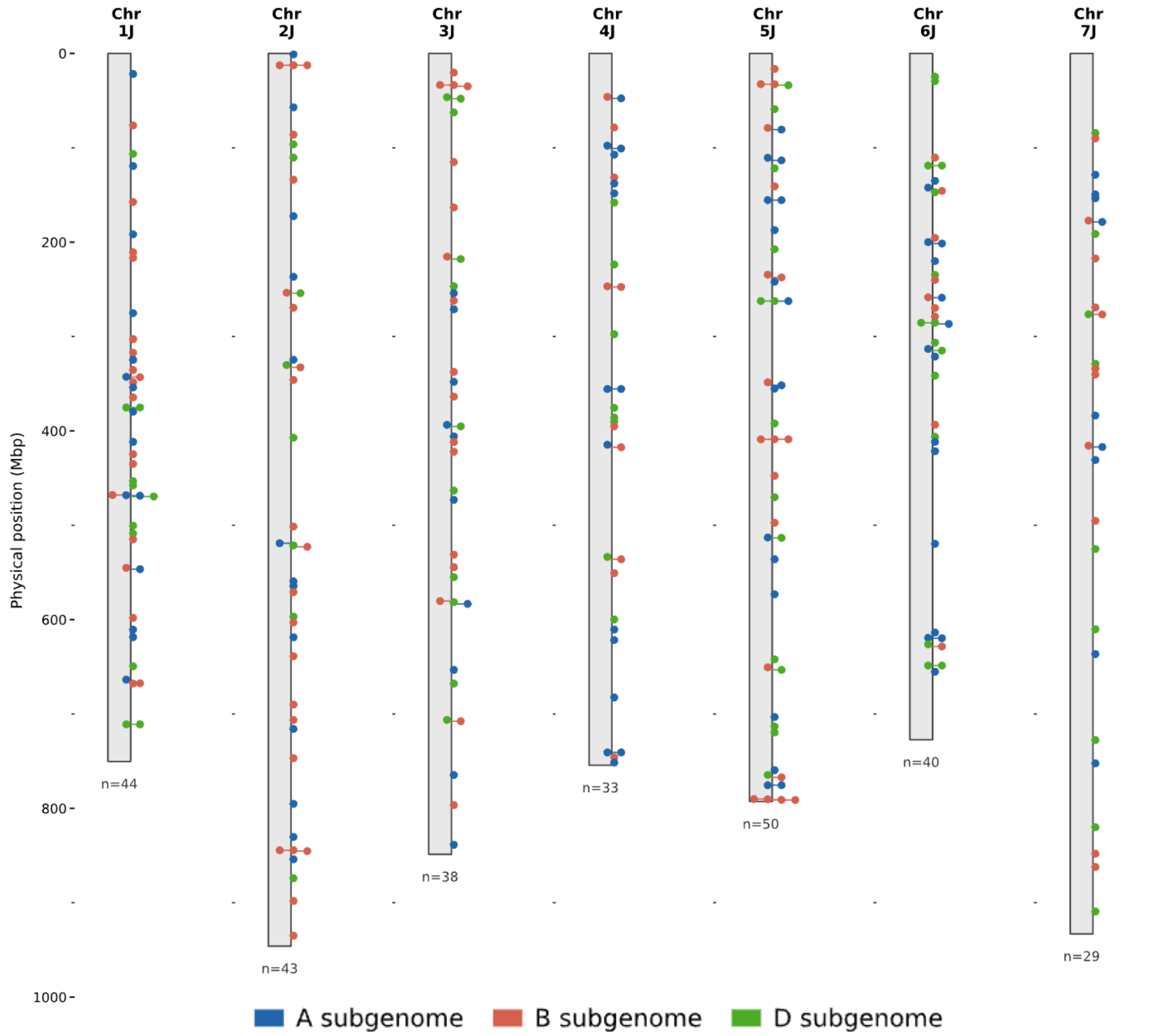


**Supplementary Figure 11. Genome-wide distribution of KASP markers across the seven *Th. bessarabicum* J-genome chromosomes.**

Each chromosome (Chr1J-Chr7J) is drawn to scale according to its physical length (Supplementary Table 3). Tick marks indicate the genomic position of each of the 277 KASP markers developed to distinguish *Th. bessarabicum* J-genome alleles from bread wheat (*T. aestivum*) alleles. Markers are coloured according to the bread wheat subgenome to which the corresponding wheat allele maps: A subgenome (blue), B subgenome (red), and D subgenome (green). The number of markers per chromosome (n) is indicated below each chromosome. Closely spaced markers are offset horizontally for clarity; their y-axis position reflects their true physical coordinate. Full marker sequences, J-genome positions, and corresponding wheat chromosomal coordinates are provided in Supplementary Table 20.

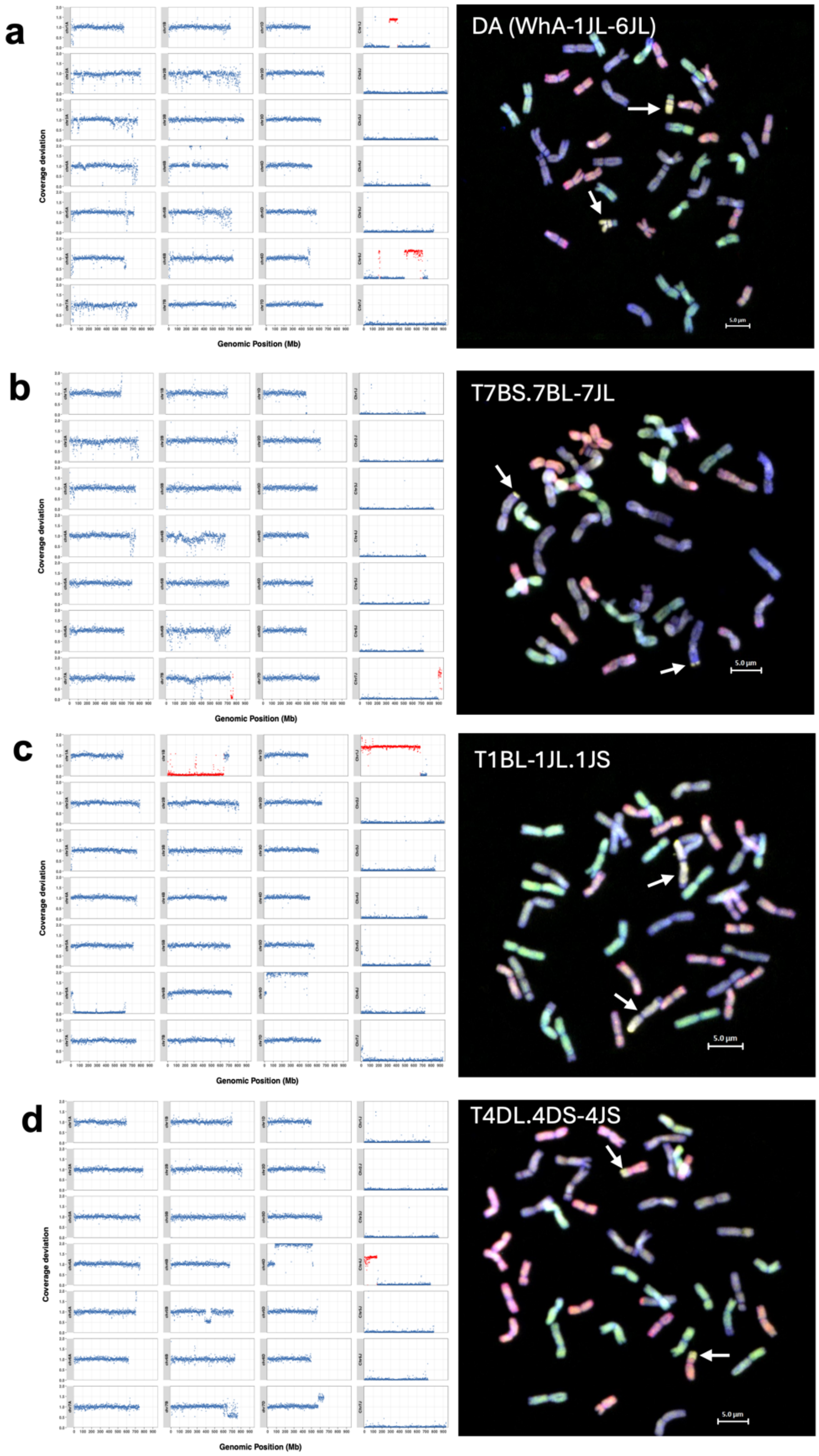


**Supplementary Figure 12. Skim-sequencing and multicolour GISH characterisation of wheat-*Th. bessarabicum* introgression lines generated at the Nottingham Wheat Research Centre (WRC).**

Skim-sequencing and multicolour GISH characterisation of four wheat-*Th. bessarabicum* introgression lines generated at the WRC. For each line, the left panel shows coverage deviation plots from the dual-reference skim-sequencing pipeline mapped simultaneously against the bread wheat RefSeq v2.1 assembly and the *Th. bessarabicum* assembly, and the right panel shows a multicolour GISH metaphase spread. In GISH images, A-genome chromosomes are shown in green, B-genome in blue-purple, D-genome in red, and *Th. bessarabicum* J-genome chromatin in yellow. White arrows indicate the introgressed J segments. In coverage deviation plots, red points indicate regions with significant deviation; values of approximately 1 indicate normal diploid coverage, approximately 1.5-2 indicate duplication (in wheat) or introgression (in J chromosomes), and approximately 0 indicate loss (in wheat) or absence of chromatin (in J chromosomes). **(a)** WRC22-Bess1 [DA(WhA-1JL-6JL)]: a novel disomic addition chromosome formed by joined segments of Chr1J and Chr6J, confirmed by simultaneous gain on both Chr1J and Chr6J in skim-seq. GISH reveals a segment of A-genome chromatin on the addition chromosome, indicative of a non-homoeologous translocation of an A-genome fragment onto the J addition chromosome. **(b)** WRC22-Bess2 [T7BS.7BL-7JL]: a small introgression from the Chr7J distal arm into wheat 7BL, confirmed by Chr7J gain (894-933 Mb) and a drop in Chr7B coverage at 736-757 Mb. **(c)** WRC22-Bess3/4 [T1BL.1JL.1JS]: a large Chr1J introgression replacing wheat chromosome 1B, confirmed by Chr1J gain (0-674 Mb) and loss of Chr1B (0-640 Mb) coverage. Sister lines WRC22-Bess3 and WRC22-Bess4 carry the same introgression thus, one representative is shown. **(d)** WRC22-Bess5 [T4DL.4DS-4JS]: an introgression from the Chr4J short arm (0-142 Mb) into a duplicated copy of wheat chromosome 4D (81-513 Mb).

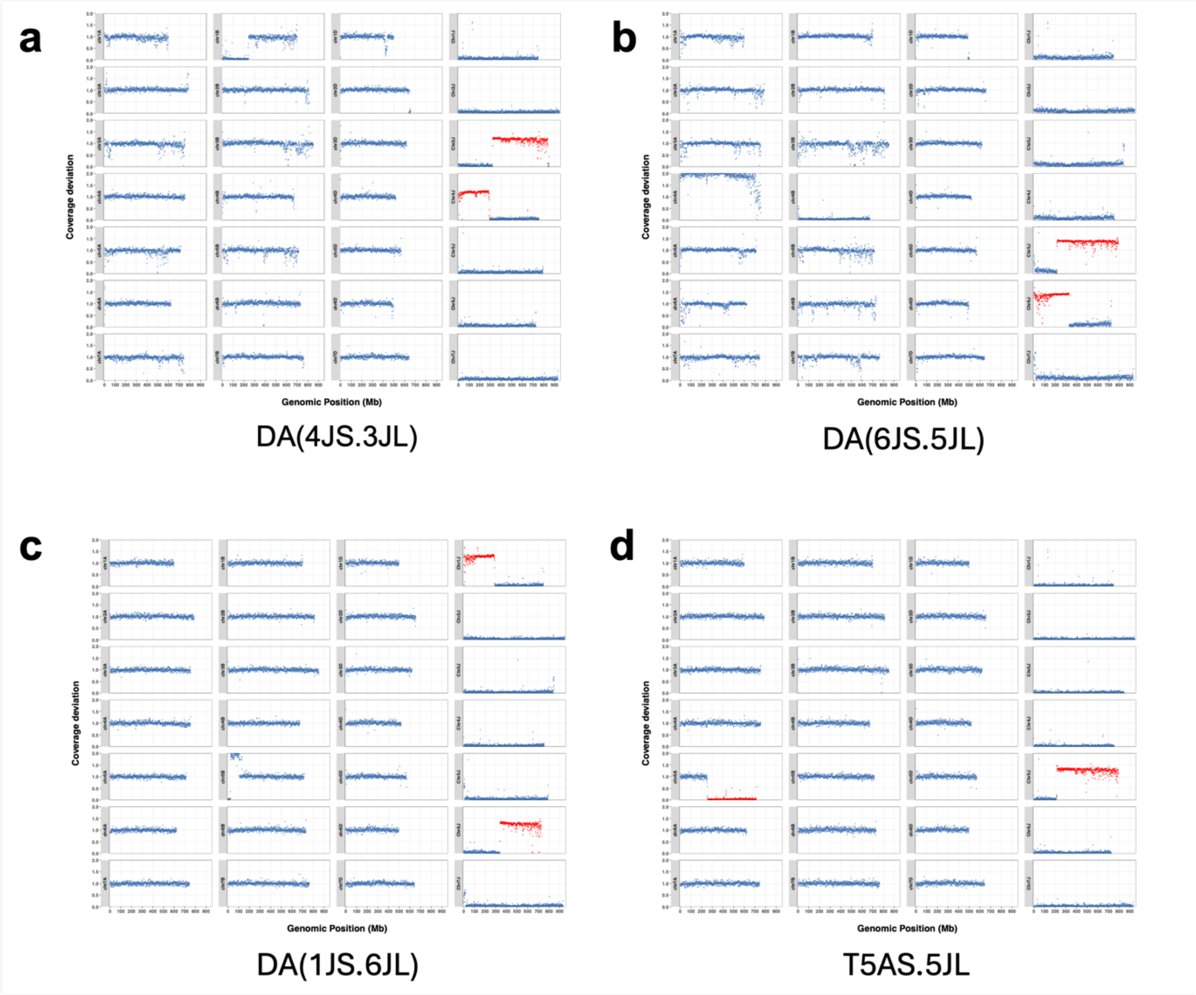


**Supplementary Figure 13. Skim-sequencing coverage deviation profiles for four wheat-*Thinopyrum bessarabicum* lines carrying centric fusions between J chromosomes and a wheat-J Robertsonian translocation.**

Coverage deviation plots from the dual-reference skim-sequencing pipeline for lines **(a)** DA(4JS.3JL), **(b)** DA(6JS.5JL), **(c)** DA(1JS.6JL) and **(d)** T5AS.5JL. For each line, the left three columns show wheat chromosome coverage (A, B and D subgenomes) and the right column shows J chromosome coverage. Red points indicate regions with significant deviation from expected coverage. The three centric fusion lines DA(4JS.3JL), DA(6JS.5JL) and DA(1JS.6JL) each show simultaneous gain on two J chromosomes, confirming their composition as centric fusions between J chromosome arms. Line T5AS.5JL shows Chr5J gain alongside a drop in chr5AS coverage, confirming a true homoeologous recombination event between Chr5J and wheat chromosome 5A. Coverage deviation values of 1 indicate normal diploid representation; values near 2 indicate duplication/introgression; values near 0 indicate absence/loss of chromatin.
